# Inferring Processes of Coevolutionary Diversification in a Community of Panamanian Strangler Figs and Associated Pollinating Wasps

**DOI:** 10.1101/490862

**Authors:** Jordan D. Satler, Edward Allen Herre, K. Charlotte Jandér, Deren A. R. Eaton, Carlos A. Machado, Tracy A. Heath, John D. Nason

## Abstract

The fig and pollinator wasp obligate mutualism is diverse (~750 described species), ecologically important, and ancient (~80-90 Ma), providing model systems for generating and testing many questions in evolution and ecology. Once thought to be a prime example of strict one-to-one cospeciation, current thinking suggests that genera of pollinator wasps co-evolve with corresponding subsections of figs, but the degree to which cospeciation or other processes contributes to the association at finer scales is unclear. Here we use genome-wide sequence data from a community of Panamanian strangler figs (*Ficus* subgenus *Urostigma*, section *Americana*) and associated fig wasp pollinators (*Pegoscapus spp.*) to infer the process of coevolutionary diversification in this obligate mutualism. Using a model-based approach adapted from the study of gene family evolution, our results indicate pervasive and ongoing host switching of pollinator wasps at this fine phylogenetic and regional scale. Although the model estimates a modest amount of cospeciation, simulations reveal this signal to be consistent with levels of co-association expected under a model of free host switching. Our findings provide an outline for testing how ecological and evolutionary processes can be modeled to evaluate the history of association of interacting lineages in a phylogenetic framework.

## 2 Introduction

Interactions among species are fundamental drivers in the diversification of life. Although many of these interactions are generalized and fleeting, some are specialized and persist over deep evolutionary timescales, playing an important role in producing coevolutionary patterns between interacting lineages (*e.g.*, Ehrlich and Raven, 1964; Thompson, 1994). For longterm associations, the ecological and evolutionary processes that have contributed to their persistence and diversification are often not well understood. For example, little is known of the types of species interactions that ultimately favor strict codiversification—where two lineages coevolve producing congruent diversification patterns—as opposed to obligately-associated lineages with relatively weak species specificity. Placing obligately-interacting lineages in an evolutionary context can inform the processes that have been important in contributing to biotic interactions, and in understanding the role interacting lineages play in generating and maintaining biodiversity.

For long-term interactions (*e.g.*, host-parasite, mutualisms), researchers often place co-evolutionary questions in a phylogenetic framework, using cophylogenetic methods to compare diversification patterns between interacting lineages, and to infer processes that may be important in contributing to these patterns. These approaches can be placed into two categories: global-fit methods and event-based methods (for review see Cruaud and Rasplus, 2016). Global-fit methods estimate the degree of congruence between two phylogenies, using either phylogenetic or genetic distance information, whereas event-based methods invoke various evolutionary events (*e.g.*, cospeciation, duplication, host switching, loss) to allow mapping of one phylogeny (*e.g.*, parasite) onto another phylogeny (*e.g.*, host). While both approaches can provide insight into coevolutionary processes, they have limitations. For example, global-fit methods measure the degree to which two data sets match but do not evaluate what processes could be contributing to any discordance. Event-based methods using the parsimony criterion (as commonly implemented) require *a priori* costs to be assigned to the various events, with the reconciliation containing the lowest total cost selected, of which there may be more than one. Both of these approaches are based on having a fixed phylogeny for each group of organisms, effectively ignoring any uncertainty in the phylogenetic estimate. This will be problematic if the input trees contain considerable—or even modest—uncertainty or do not adequately capture the diversification history. Importantly, these approaches do not directly measure the contributions of the coevolutionary processes that can generate the observed data in a probabilistic framework. As multiple processes can generate data with similar patterns, it is necessary to use a model-based approach where the relative contribution of these processes can be evaluated in a statistical framework. Thus, insight into the evolutionary history of coevolutionary associations will benefit from a shift in how cophylogenetic questions are framed and tested.

Obligate mutualisms provide important systems to understand the processes that contribute to the maintenance and diversification of interacting lineages. The mutualism between fig trees and their pollinating fig wasps is one of the best-known examples of an obligate mutualism. Figs are an ecologically and morphologically diverse genus (*Ficus*) of over 750 described species with a pantropical distribution. They are considered keystone species in tropical ecosystems because their generally aseasonal production of large fruit crops contributes heavily to the diets of many vertebrates, especially during seasons in which fruiting activity of other tree species is low (Terborgh, 1986; Lambert and Marshall, 1991; Kissling et al., 2007). Each species of fig is pollinated by a specialized fig wasp, with both members of this mutualism solely dependent on the other for reproduction and survival—the fig requires the fig wasp for pollination services, and the fig wasp requires the fig as a nursery for larval development (Wiebes, 1979). This obligate association has been maintained for upwards of 90 million years (Machado et al., 2001, 2005; Rønsted et al., 2005).

Decades of research have provided a foundation of knowledge into the ecology and evolution of this specialized interaction, providing insight into mechanisms that maintain the mutualism, as well as the factors that structure coevolutionary patterns. Although the long-held paradigm has been a one-to-one relationship between individual fig and fig wasp species, analyses of molecular data sets have revealed examples of multiple pollinator species associated with a single fig species as well as multiple fig species that share a single pollinator species (*e.g.*, Molbo et al., 2003; Haine et al., 2006; Marussich and Machado, 2007; Darwell et al., 2014; Yang et al., 2015; Wang et al., 2016). While a signal of cospeciation is generally recovered between sections of figs and genera of fig wasps (*e.g.*, Rønsted et al., 2005; Cruaud et al., 2012), phylogenetic data have often revealed discordant diversification patterns within these groups, requiring additional evolutionary processes (*e.g.*, host switching) to explain observed deviations from the expected one-to-one pattern of cospeciation (Machado et al., 2005; Jackson et al., 2008).

To date, these results have been based on Sanger sequencing of one or few genes, often resulting in poorly resolved estimates of phylogenetic relationships. With advances in sequencing technologies and the ability to gather hundreds to thousands of loci for non-model organisms, large genomic data sets can be used to decrease sampling error and generate better phylogenetic estimates with greater confidence. In addition, it is important to account for incomplete lineage sorting in phylogenetic methods, as this biological process can introduce substantial error when unaccounted for in phylogenetic inference (*e.g.*, Kubatko and Degnan, 2007). Thus, robust phylogenies coupled with model-based approaches can now be used to rigorously test how various processes have contributed to observed patterns of interspecific association, providing insight into the coevolution and diversification of host-symbiont associations, including mutualism, such as figs and fig wasps, and antagonism, such as host-parasite interactions.

Here we use genome-wide sequence data to infer the processes governing coevolutionary diversification in a community of Panamanian strangler figs and their pollinating fig wasps. We gather restriction-site associated loci (RAD) from host strangler figs and ultraconserved element loci (UCE) from their associated pollinating wasps, and estimate phylogenetic relationships to test how coevolutionary processes have contributed to the evolution of this mutualism. Using a probabilistic approach, we adapt a duplication-transfer-loss (DTL) model of gene family evolution (Szöllősi et al., 2012) to quantify the contribution of fundamental ecological and evolutionary processes to this mutualism, including wasp speciation, host switching, and wasp extinction. Already accounting for uncertainty in the fig wasp species tree distribution, we extend the model implementation to account for phylogenetic uncertainty in the host figs, and introduce simulations of null hypotheses to aid biological interpretation of results. In sum, we present a framework for addressing coevolutionary questions using a model-based method, and demonstrate the benefits of utilizing this approach when inferring the evolutionary history of long-term interactions.

## 3 Materials & Methods

### 3.1 Sampling, sequencing, and phylogenetics of figs

More than 120 species of strangler figs (family Moraceae, *Ficus* subgenus *Urostigma*, section *Americana*) have been described from the Neotropics. We sampled 31 individuals from 11 species of strangler figs (family Moraceae, *Ficus* subgenus *Urostigma,* section *Americana*) co-occurring in central Panama, centered around the Barro Colorado Island Nature Monument (Table 1). These 11 fig species include two pairs that share pollinator species, and another fig species that hosts two pollinator wasp species (Table 1). The advantage of focusing on this particular set of species is that this Panama fig community has been extensively studied over two decades providing long-term ecological and genetic data (*e.g.*, Herre et al., 2008; Nason et al., 1996; Machado et al., 2005; Molbo et al., 2003). Genomic DNA was extracted using a modified CTAB protocol (Doyle and Doyle, 1987) and sent to Floragenex Inc. (Eugene, OR, USA) for library preparation using the PstI restriction enzyme and the traditional RAD protocol (Baird et al., 2008). The resulting library was sequenced on two lanes of an Illumina HiSeq 3000 using 100 base pair single-end sequencing. Raw sequencing reads were processed in ipyrad v0.6.20 (Eaton, 2014) to generate alleles, loci, and SNPs. Sequences were first demultiplexed, not allowing any mismatches with the barcode, and then filtered for Illumina adapter contamination. Base calls with a Phred score below 30 were replaced with an N; up to 5 Ns were allowed in any given sequence, with sequences above that threshold removed from the data set. A clustering threshold of 85% was used to assemble reads into loci. When clustering loci among samples, we allowed up to four individuals to be heterozygous for a shared site, up to 20 SNPs in a locus, and up to 8 indels in a locus. All loci that contained at least four individuals were retained in the complete data set. We furthered filtered this pool to those loci that contained at least one individual per species, then subsampled a single SNP per locus at random for analysis.

Two approaches were used to generate species-level phylogenetic estimates of the sampled fig trees. As both analyses use the multispecies coalescent model, individuals were assigned to their respective species *a priori*. First, SNAPP v1.3 (Bryant et al., 2012) in BEAST v2.4.7 (Bouckaert et al., 2014) was used to estimate a posterior distribution of species trees from biallelic SNP data under the multispecies coalescent model. SNPs were recoded as “0, 1, 2”, with heterozygotes as “1”, major allele at “0”, and minor allele at “2”. Major and minor allele frequencies were calculated in BEAUti informing priors on forward and reverse mutations; all other settings were left at default. Analyses were run for 2 million generations, sampling every 200 generations, resulting in 10,000 Markov chain Monte Carlo (MCMC) samples from the posterior distribution. To ensure consistency across runs, two independent Markov chains were run. Resulting log files were imported into Tracer v1.6.0 (Rambaut et al., 2018) to assess non-convergence. In addition, a species tree was estimated with SVDquartets (Chifman and Kubatko, 2014) as implemented in PAUP* v4.0a163 (Swofford, 2003). SVDquartets uses site patterns in the nucleotide data to estimate a species tree under the multispecies coalescent model. All quartets were evaluated exhaustively with 100 bootstrap replicates run to evaluate nodal support values.

**Table 1.** Sampling and ecological associations of figs and fig wasps. Species, number of individuals sequenced (N), and fig and fig wasp associations are shown. An asterisk (*) indicates pollinating wasp species shared by two host figs.

| Fig species | N | Fig wasp species | N |
| --- | --- | --- | --- |
| *Ficus citrifolia* | 2 | *Pegoscapus tonduzi* | 5 |
| *Ficus costaricana* | 2 | *Pegoscapus estherae* | 6 |
| *Ficus nymphaeifolia* | 3 | *Pegoscapus piceipes* | 4 |
| *Ficus paraensis* | 3 | *Pegoscapus herrei* | 7 |
| *Ficus near trigonata* | 3 | *Pegoscapus lopesi* | 10 |
| *Ficus trigonata* | 3 | *Pegoscapus grandii* | 8 |
| *Ficus obtusifolia* | 3 | *Pegoscapus hoffmeyerii* A | 2 |
|  |  | *Pegoscapus hoffmeyerii* B | 3 |
| *Ficus perforata* | 3 | *Pegoscapus insularis* * | 15 |
| *Ficus colubrinae* | 3 |  |  |
| *Ficus popenoei* | 3 | *Pegoscapus gemellus* B | 9 |
| *Ficus bullenei* | 3 | *Pegoscapus gemellus* C | 10 |
|  |  | *Pegoscapus gemellus* A* | 3 |

### 3.2 Sampling, sequencing, and phylogenetics of fig wasps

We sampled 82 individuals from 12 lineages of pollinating fig wasps (family Agaonidae, genus *Pegoscapus*) associated with the 11 focal strangler fig species (Table 1). Wasps were sampled from the fig tree species sequenced above and allowed to emerge from mature fig fruit in the lab, where they were then stored in 95% EtOH or RNALater for DNA collection and analysis. A single wasp was sequenced per fig fruit to ensure independence among samples. Genomic DNA was extracted with a Qiagen DNeasy Kit (Qiagen Inc., Valencia CA, USA), and amplified using a Qiagen REPLI-g Mini Kit to increase total DNA yield. On average, 10-20 ng of DNA was used for whole genome amplification. Samples were standardized and sent to RAPiD Genomics (Gainsville, FL, USA) for library preparation and sequencing. The hymenopteran probe v1 set of Faircloth et al. (2015) was used to target 1510 ultraconserved element (UCE) loci. UCE loci have been shown to be informative phylogenetic markers across a broad group of animals, including wasps (*e.g.*, McCormack et al., 2012; Crawford et al., 2012; Branstetter et al., 2017; Starrett et al., 2017). Following library construction, samples were sequenced on half a lane of an Illumina HiSeq 2000 using 150 base pair paired-end sequencing. Phyluce v1.5 (Faircloth, 2015) was used to process raw sequence reads and generate loci for each individual. Briefly, sequence reads were cleaned with Trimmomatic (Bolger et al., 2014), and assembled into contigs with Trinity v2.0.6 grabherretal2011. Contigs were then aligned to the hymenopteran v1 UCE locus set to filter non-specific sequences. Loci were then aligned with MAFFT (Katoh and Standley, 2013), edge trimmed with trimAl (Capella-Gutiérrez et al., 2009), and ambiguously aligned internal sites removed with Gblocks castresana2000. We retained loci sampled for a minimum of 70% of individuals in the final data set, resulting in 550 UCE loci.

Two approaches were used to generate phylogenetic estimates of the pollinator wasps. First, a species tree was estimated using StarBEAST2 v0.14.0 (Heled and Drummond, 2010) in BEAST v2.4.7. StarBEAST2 coestimates the posterior distributions of gene trees and species trees under the multispecies coalescent model. This approach accounts for uncertainty in the tree topologies, branching times, and other model parameters using MCMC in a Bayesian inference framework. Due to the computational challenges of applying this method, loci were first filtered to those that had *at least* one sequence per OTU per locus, resulting in a pool of 353 UCE loci. We then randomly subsampled *without replacement* 50 loci for analysis. Preliminary analyses containing more loci failed to reach stationarity. In addition, a maximum of three alleles per species per locus was used to reduce computational burden, as this is sufficient for species tree estimation (Hird et al., 2010). For species at or below this allelic threshold for a given locus, all alleles were used, otherwise three alleles were randomly subsampled *with replacement* for analysis. We used a birth-death model as a prior on the species tree, an HKY + I + Γ substitution model, and a strict clock for each locus. Analyses were run for two billion generations, sampled every 200,000 generations, yielding 10,000 samples from the target distribution. Two independent analyses were run and non-convergence was diagnosed using Tracer. We also estimated a species tree using SVDquartets, with all 550 UCE loci included for analysis. In addition to estimating a species tree with individuals assigned to species (*i.e.*, SVDQ_*ST*_), we estimated a phylogeny using the individuals as tips in the tree (*i.e.*, a lineage tree; SVDQ_*LT*_). The lineage tree (SVDQ_*LT*_) was primarily used to confirm species association. For both analyses, quartets were evaluated exhaustively and 100 bootstrap replicates were run to evaluate nodal support values.

### 3.3 nferring coevolutionary processes in a model-based framework

We are interested in inferring the evolutionary and ecological processes responsible for producing the cophylogenetic patterns between interacting taxa, such as hosts and parasites, and strangler figs and their pollinating wasps. Using a model-based approach, we can evaluate the relative contribution of different parameters in a statistical framework informing what processes are important for generating our observed data. Here we adapt a model of gene family evolution to test how various processes may have contributed to estimated phylogenetic patterns. The duplication-transfer-loss (DTL) model of Szöllősi et al. (2012) estimates the maximum likelihood rates of gene duplication (D), horizontal gene transfer (T), and gene loss (L) of a gene family, given an ultrametric species tree with nodes ordered by speciation time. Then, based on the DTL rates, the probabilities of these events along the branches of a species tree can be calculated (Szöllősi et al., 2013b). Following likelihood calculation and maximization of parameters given the data, reconciled gene trees can be sampled from the species tree and event probabilities given stochastic backtracking along the species tree (Szöllősi et al., 2013a). After sampling a set of reconciled gene trees, the average number of duplication, transfer, and loss events can be summarized providing an estimate as to the average number of events for a gene family given a species tree.

The DTL model lends itself well to cophylogenetic questions because the model parameters estimated are analogous to those in our system—a host phylogeny represents the species tree and a symbiont phylogeny represents the gene-family tree. For example, modeled processes that may contribute to the coevolution of the fig (host) and pollinating fig wasp (symbiont) include duplication (*i.e.*, wasp speciation, where a wasp lineage speciates on a fig resulting in two pollinating species on one host), transfer (*i.e.*, host switching, where a wasp lineage colonizes a new fig species host), and loss (*i.e.*, wasp extinction, where a fig speciates but the associated wasp fails to speciate and tracks one fig daughter lineage, or the wasp lineage goes extinct). The sampled reconciled tree set provides average values of these three events in addition to counts of cospeciation (*i.e.*, the number of times a speciation event in the host lineage co-occurred with speciation in its symbiont). Thus, this method estimates how cospeciation, host switching, fig wasp speciation, and fig wasp extinction contribute to the coevolutionary diversification patterns in interacting lineages.

We applied the DTL model of gene family evolution for estimating the evolutionary processes contributing to cophylogenetic patterns as implemented in the dated version of the program ALEml (https://github.com/ssolo/ALE). ALEml takes as input an ultrametric species tree, with nodes ordered by speciation time, and a posterior distribution of gene family trees, accounting for uncertainty in the gene family tree estimates. In our application of the model, we treat the fig species tree as the host species tree, and the posterior distribution of fig wasp species trees as the gene family trees, explicitly accounting for phylogenetic uncertainty in the fig wasps. For the two cases where a single wasp species pollinates two different fig tree species, we split the wasp tip into two sister tips for all fig wasp species trees sampled in the posterior distribution from StarBEAST2. This was applied to *P. insularis* (a wasp species that pollinates both *F. colubrinae* and *F. perforata*) and *P. gemellus A* (a wasp species that pollinates both *F. popenoei* and *F. bullenei*). ALEml was then used to estimate maximum likelihood rates of wasp speciation (*i.e.*, duplication), host switching (*i.e.*, transfer), and wasp extinction (*i.e.*, loss), generating a set of 1,000 reconciled fig wasp species trees given the host species tree and probability of events along the branches. Averaging over this set of sampled reconciled fig wasp species trees provided average number of wasp speciation, host switching, wasp extinction, and cospeciation events. We extended the implementation of the method to also account for phylogenetic uncertainty in the fig species tree by thinning the post-burnin SNAPP posterior distribution to 1,000 species trees, and running the analysis across all 1,000 trees sequentially. Results were then averaged across these 1,000 replicates.

### 3.4 Simulations

Simulations were used to evaluate the behavior and sensitivity of ALEml, and to contextualize empirical estimates by generating null distributions with which to compare estimated values. Specifically, trees were simulated with no explicit assumption of association between the tips of the simulated trees and the empirical host fig tree, providing an expectation of parameter values when there is no explicit history of cospeciation. These simulations approximate expectations under a model of free host switching, providing a null model with which to compare empirical parameter estimates. We simulated 1,000 trees under a birth-death process (λ=1.0, *μ*=0.9) in the R package TreeSim v2.3 (Stadler, 2011). Tree distributions were simulated with the sampling properties of the fig wasps (*i.e.*, three fig tip species were randomly assigned two simulated tips, all other fig tips were randomly assigned a single tip). For the ALEml analyses, the fig maximum-clade credibility (MCC) tree summarized from the SNAPP analysis was used, keeping the host tree constant, with one simulated tree used at a time until all 1,000 trees in the set were independently analyzed. Using a single tree to represent the fig wasps removes phylogenetic uncertainty, but is sufficient for the purposes of the simulations. Parameter estimates across all replicates were then summarized as null distributions for the model parameters.

### 3.5 Describing cophylogenetic patterns using traditional approaches

In addition to our application of the gene-family evolution model to analyze codiversification, two traditional approaches were used to understand this mutualism. First, the degree of congruence between the fig and fig wasp phylogenies was assessed in the R package PACo v0.3.2 (Balbuena et al., 2013; Hutchinson et al., 2017). PACo is a global-fit method that uses Procrustean superimposition to evaluate if two phylogenies are more similar than expected by chance, suggesting a shared evolutionary history. An eigenvalue correction of “lingoes”—a constant added to correct for negative eigenvalues—was used when transforming the phylogenetic distance matrices into principal coordinates. Rather than superimpose the fig wasp phylogeny on the fig phylogeny (as would be the case in a host and parasite system), both phylogenies were standardized to allow for the best-fit superimposition independent of both phylogenies. We used 10,000 permutations to assess significance for the analysis. Second, the parsimony event-based method Jane v4.0 (Conow et al., 2010) was used to estimate the number of events required to explain discordance in the two phylogenies. Jane takes as input two phylogenies (one from either group) and reconciles the two trees by minimizing the number of event costs. Costs are applied *a priori*; default values were used with cospeciation = 0, host switching = 2, duplication = 1, loss = 1, and failure to diverge = 1. Using the parsimony criterion, the optimal reconciliation is the one that minimizes the total cost. Following the reconciliation process, statistical support was generated by drawing 1.0 random pollinator trees from a Yule model (default beta value of −1.0) to generate a null distribution of event costs. For PACo and Jane, MCC trees from SNAPP (figs) and StarBEAST2 (figwasps) were used as input data.

## 4 Results

### 4.1 Sampling, sequencing, and phylogenetics of figs

We generated 228,004,993 raw reads for the 31 sampled fig trees, with an average of 7,355,000 reads per individual (± 9,704,999). Following data processing in ipyrad, individuals had on average 18,444 loci (± 9,023). Requiring at least one individual to be sequenced per species per locus for phylogenetic analysis, 2529 unlinked biallelic SNPs were used to estimate a species tree for the figs. Phylogenetic results were generally consistent between SNAAP and SVDquartets. With SNAPP, species fell into two main clades, with generally good support throughout the tree although some nodes reflected greater uncertainty (Figure 1). For the figs that share a fig wasp species, drastically different patterns emerge. *Ficus bullenei* and *F. popenoei* are sister species in the phylogeny. In contrast, while *F. perforata* and *F. colubrinae* fall in the same clade, they are not closely related. Similar interrelationships are recovered with SVDQ_*ST*_, with varying levels of support across the tree (Figure S1). One difference from SNAPP, however, is *F. perforata* and *F. colubrinae* are sister taxa in the SVDQ_*ST*_, although this is weakly supported (BS = 67).

**Figure 1:**
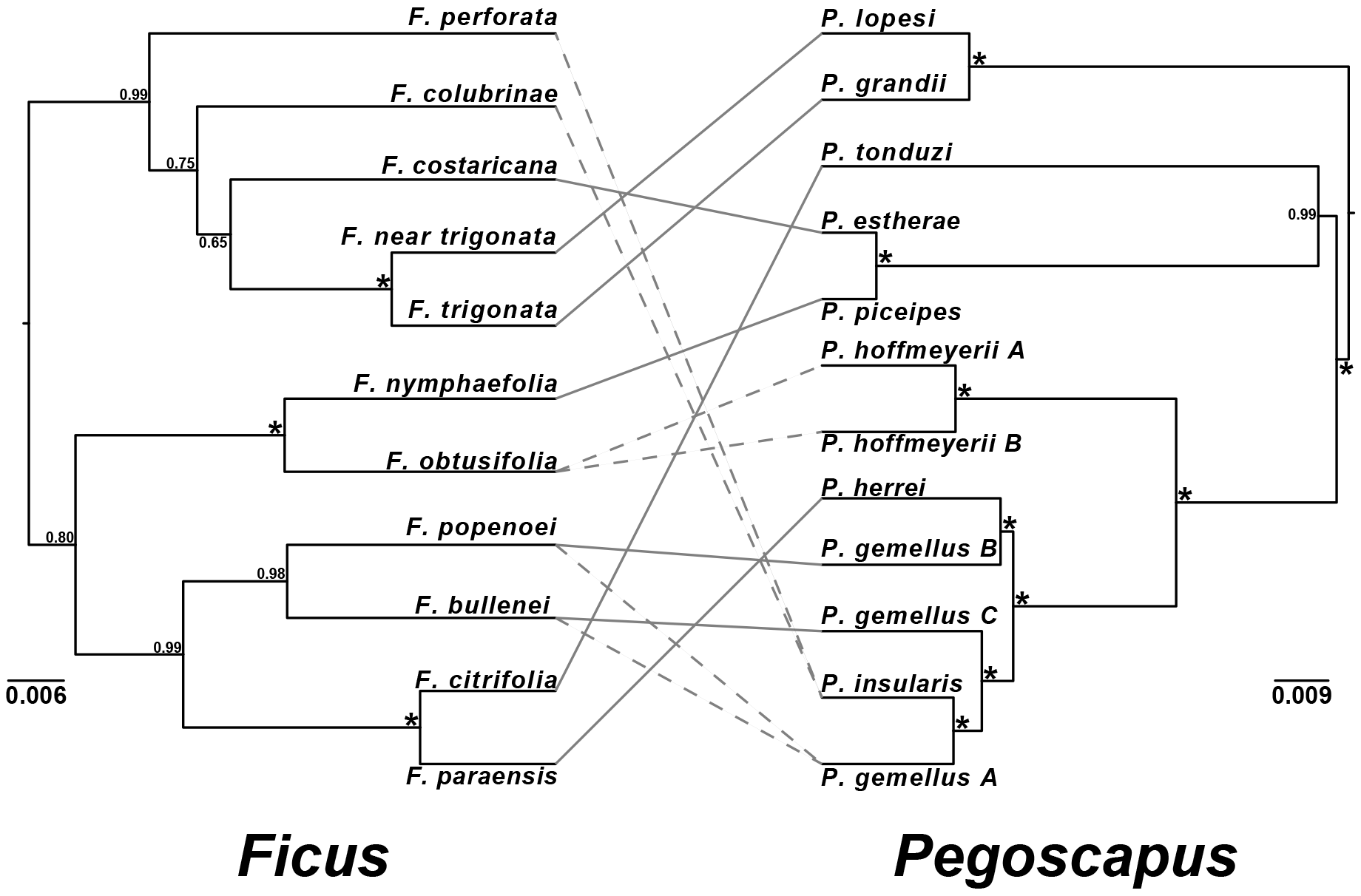
Tanglegram representing evolutionary relationships and associations of 11 species of Panamanian strangler figs (*Ficus*) and and their respective pollinating fig wasps (*Pegoscapus*). The species tree MCC estimates of the figs (SNAPP) and fig wasps (StarBEAST2) are shown with nodal support values; posterior probabilities of 1.0 are represented with an asterisk. Broken lines in associations represent when one-to-one relationship breaks down.

### 4.2 Sampling, sequencing, and phylogenetics of fig wasps

We generated 164,703,731 raw reads for the 82 sampled fig pollinating wasp individuals, resulting in an average of 2,008,582 reads per individual (± 577,848). Following processing in phyluce, a pool of 1052 total UCE loci was recovered, of which 1017 contained at least three or more individuals per locus. On average, there were 628 loci per individual (± 162.60). Requiring a minimum of 70% taxon coverage to retain a locus, the final data set consisted of 550 loci. Loci in this data set had an average size of 534 bp (± 137.62), ranging from 122 bp to 1,072 bp, and were present in an average of 64 individuals (± 4.84). The fig wasp species trees resulting from StarBEAST2 and SVDQ_*ST*_ had generally congruent topologies. *Pegoscapus* wasps are grouped into three main clades, with all nodes strongly supported (Figure 1). In the two cases where single wasp species are associated as pollinators to two host species, the wasp species are sister in the phylogeny (*P. gemellus_A* (Hosts: *F. bullenei* and *F. popenoei*) sister to *P. insularis* (Hosts: *F. colubrinae* and *F. perforata*)). SVDQ_*ST*_ shows the same general pattern as the StarBEAST2 trees, with one minor exception—*P. tonduzi* (Host: *F. citrifolia*) is sister to *P. piceipes* (Host: *F. nymphaefolia*), with *P. estherae* (Host: *F. costaricana*) sister to them (Figure S2). This node, however, has a bootstrap value of 50, showing little support for this discordant pattern. All other nodes have 100 bootstrap support. SVDQ_*LT*_ shows the same branching pattern as SVDQ_*ST*_, and demonstrates close genetic relatedness among wasp individuals as determined by host fig species (Figure S3), supporting our placement of individuals within species for species tree analysis.

### 4.3 Inferring coevolutionary processes in a model-based framework

Integrating across phylogenetic uncertainty in the sample fig and fig wasp community, our application of a DTL model for cophylogenetics estimated mean maximum likelihood rates of 2.59 (± 0.83) for host switching, 1.17 (± 0.42) for wasp extinction, and 0.06 (± 0.04) for wasp speciation (Figure 2). Given these rates and the probabilities of events along the branches of the host fig trees, the reconciled fig wasp tree set recovers a mean average of 11.56 (± 1.32) host switches, 5.10 (± 1.55) cospeciations, 3.45 (± 0.91) wasp extinctions, and 0.27 (± 0.12) wasp speciations (Figure 3). These results highlight the preponderant role host switching has played in producing the coevolutionary patterns between this group of strangler figs and their associated pollinating wasps, but also suggest a role for cospeciation and wasp extinction during the evolution of these co-occurring mutualisms.

**Figure 2:**
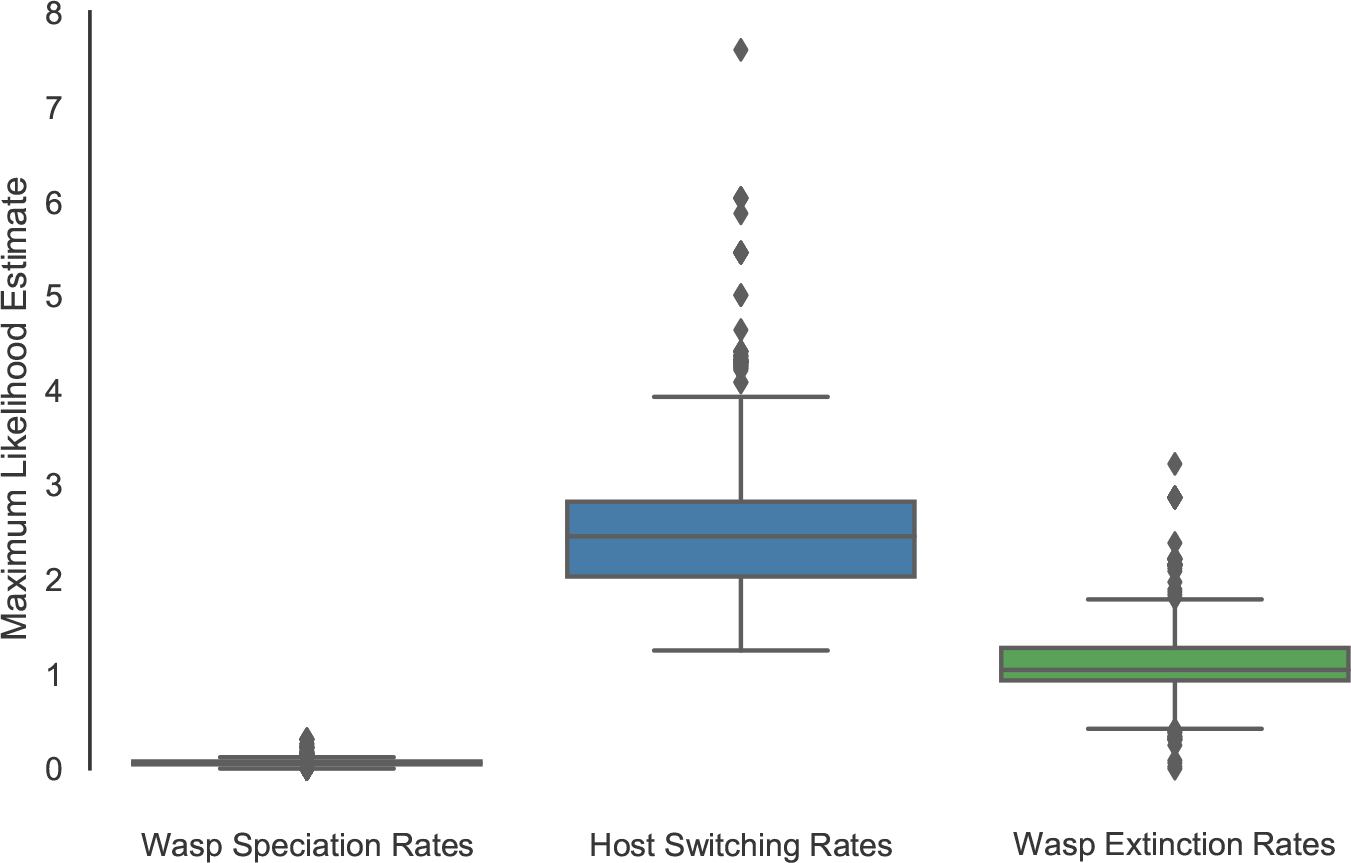
Maximum likelihood estimates for the rates of the three parameters estimated in ALEml. Results are averaged across 1,000 trees drawn from the posterior distribution of species trees sampled from the analysis (SNAPP) of the fig data.

Because there is no explicit association between the simulated trees and empirical trees, the simulations provide a context for evaluating empirical parameter estimates against expectations of no cospeciation. When we compare the average number of wasp speciations and wasp extinctions events, the empirical estimates do not significantly deviate from the null distributions (Figure 4). For the average number of host switches and cospeciations, once again, the empirical estimates do not significantly deviate from the null distributions, although both of these estimates are close to the tails of the distributions, with 15% of simulations at or below the empirical estimate for host switching, and 11% at or above the empirical estimate for cospeciation. This same general pattern is seen with the maximum likelihood rates when placed in the context of the simulations (Figure S4). These results show the empirical parameter estimates to be consistent with a model that approximates free host switching.

**Figure 3:**
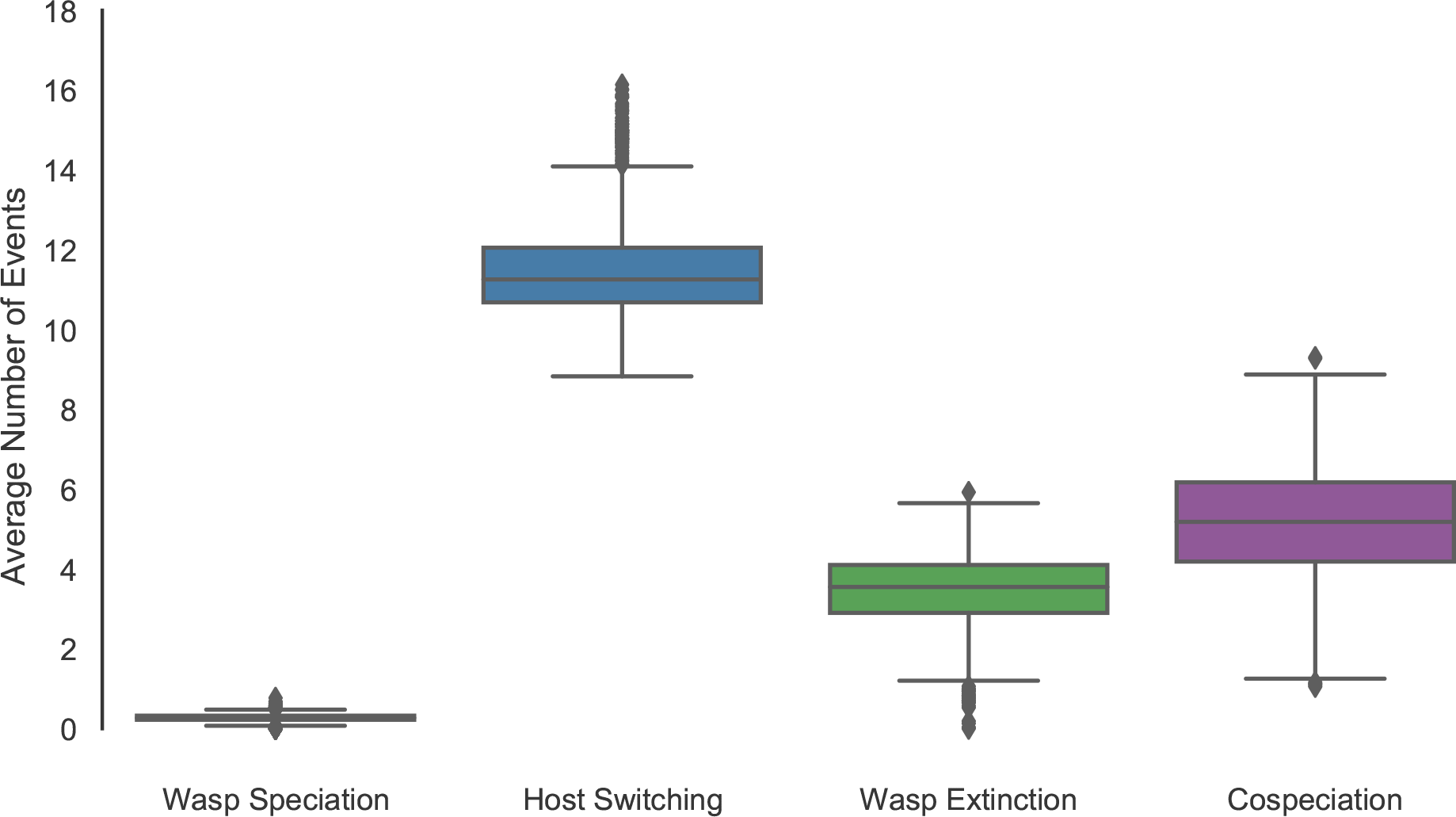
Average number of events estimated from ALEml. Results are averaged across 1,000 trees drawn from the posterior distribution of species trees sampled from the analysis (SNAPP) of the fig data.

### 4.4 Describing cophylogenetic patterns using PACo and Jane

PACo recovers a significant cophylogenetic signal between the figs and the fig wasps (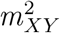 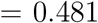, *P* = 0.009, *n* = 10,000). This demonstrates the degree of congruence between the associations is greater than expected based on chance (Figure 5). This result, however, is driven by the *F. trigonata* + *F. triangle* contribution (Figure 5A). If these two fig species (and associated pollinating wasps) are removed from the analysis, the interaction is no longer statistically significant (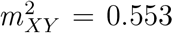, *P* = 0.11, *n* = 10,000), demonstrating their importance to the congruent cophylogenetic signal. Jane does not recover a significant association between the figs and fig wasps. There were forty-eight equally parsimonious reconciliations with a total event cost of 20. Forty-two reconciliations contained five cospeciations, three fig wasp speciations, three host switches, nine losses, and two failures to diverge (Figure 6). The other six reconciliations contained four cospeciations, two fig wasp speciations, five host switches, six losses, and two failures to diverge. When assessing significance of this cost compared to a null distribution, 25.2% of random trees contained a cost equal to or below this value, with a mean cost number of 23.33 (± 4.31).

**Figure 4:**
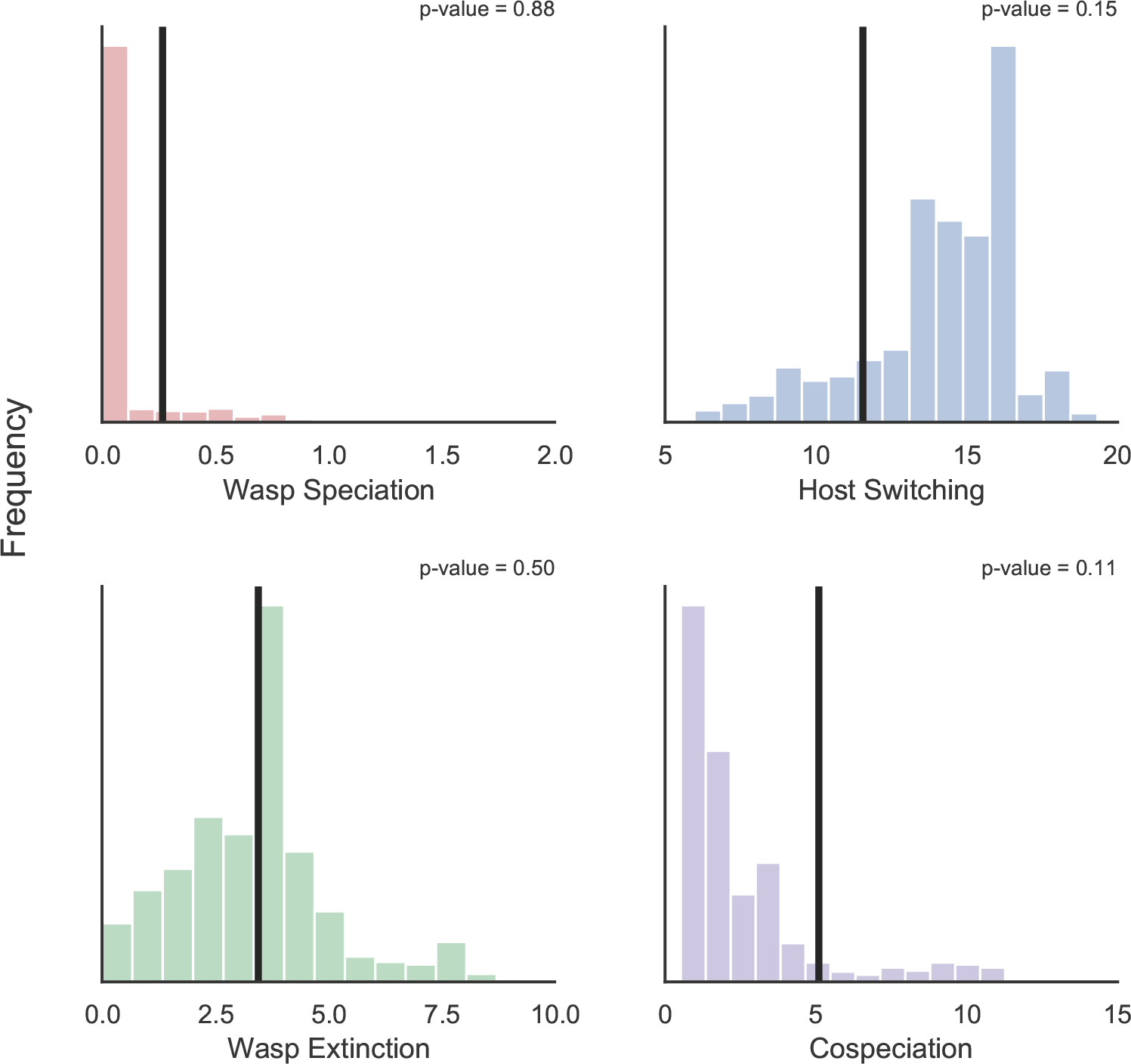
Number of events when using the fig MCC species tree and a simulated fig wasp species tree. Null distributions are based on 953 simulated fig wasp species trees (some ALEml analyses returned an error and were subsequently skipped). The vertical line represents the estimate from the empirical data. For wasp speciation, wasp extinction, and host switching, p-values represent the proportion of simulations that produced an estimate equal to or below the empirical value. For cospeciation, the p-value (0.11) represents the proportion of simulations that produced an estimate equal to or above the empirical value.

## 5 Discussion

Estimating the relative frequency of ecological and evolutionary processes that contribute to diversification patterns in interacting lineages is a fundamental goal of evolutionary biology. Here we gathered genome-wide sequence data for a community of Panamanian strangler figs and pollinating wasps, and using a model-based framework, demonstrated that host switching has been an important process contributing to the coevolution of this mutualism. These results provide direct evidence that host switching is perhaps the most dominant process ongoing at this phylogenetic level, and within the Panamanian community, provide support for an evolutionary history in which fig wasps have been able to shift to different host figs through time.

**Figure 5:**
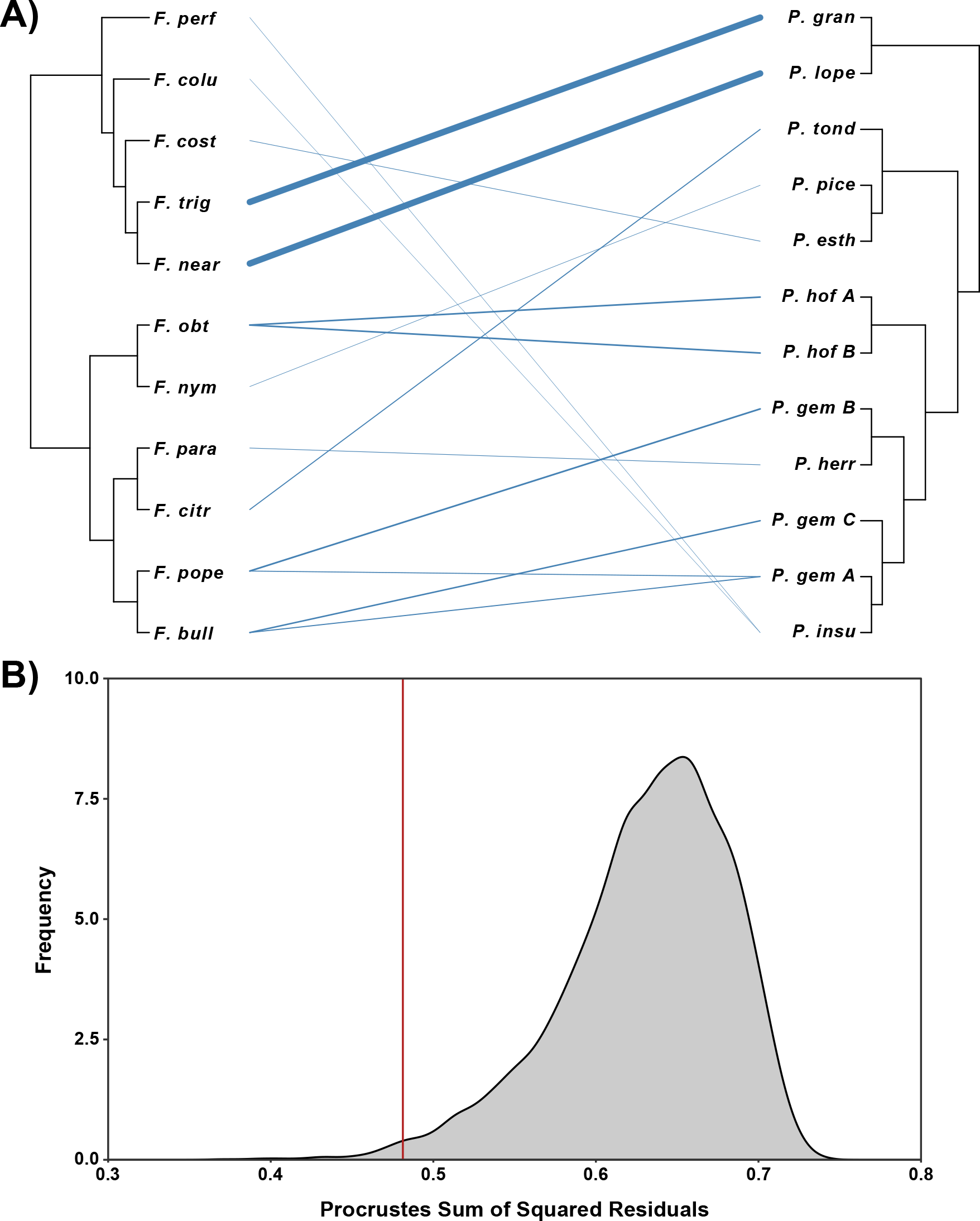
Results from PACo. (A) Associations between host figs and pollinating wasps, with weight of line showing the contribution of each interaction to the global-fit score. (B) A significant association is recovered for this mutualism. As the *F. trigonata* + *F. near trigonata* relationship contributes the most to this interaction, if we remove these two taxa (and their associated wasps), the global association is no longer significant.

**Figure 6:**
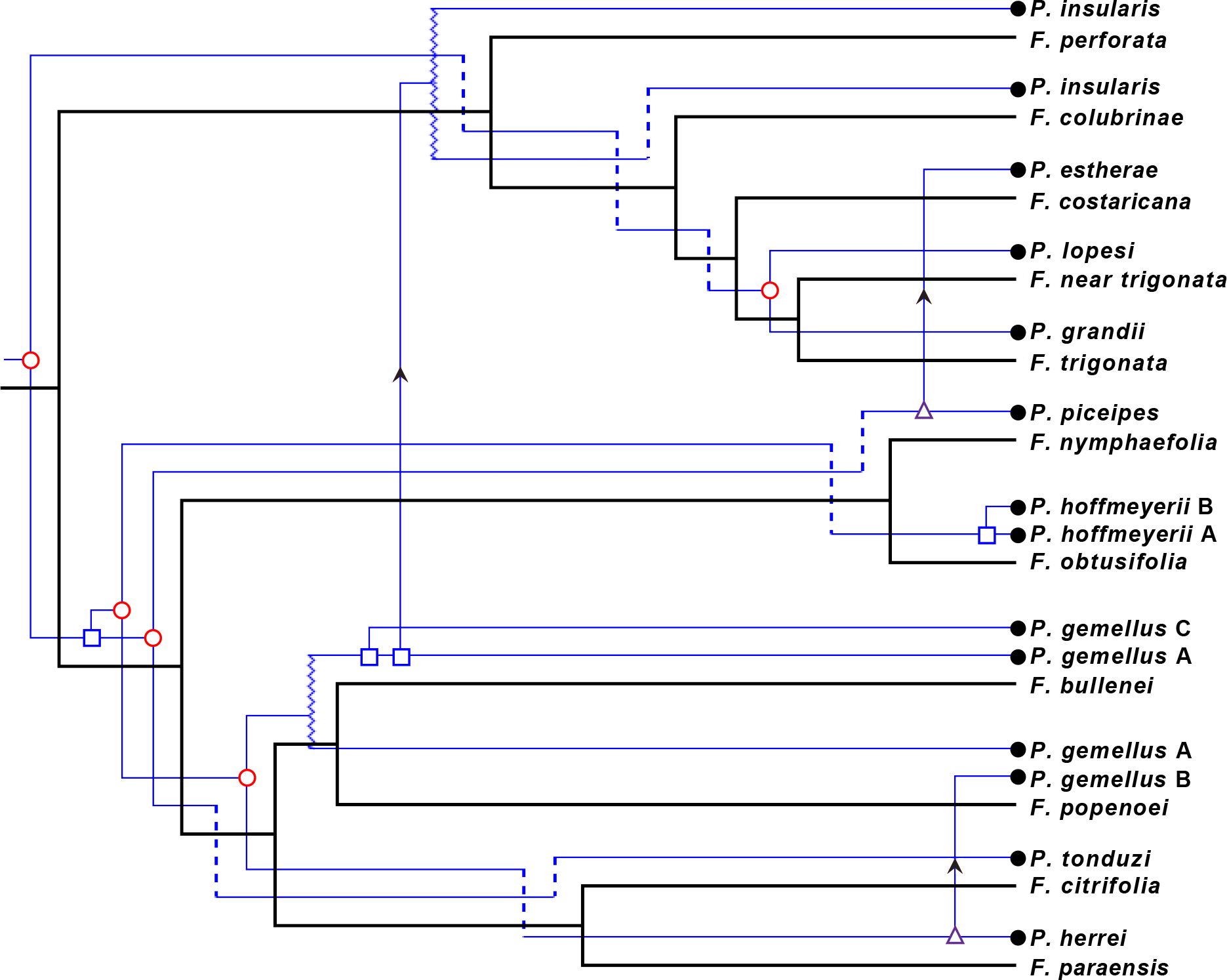
One of the equally most parsimonious reconciliations inferred by Jane. The black tree represents the host fig phylogeny, and the blue tree represents the fig wasp phylogeny with mapped events that produce the best reconciliation between these two trees. Red circles represent cospeciation, blue squares represent fig wasp speciation, purple triangles represent fig wasp speciation followed by a host switch, broken lines represent loss, and jagged lines represent failure to diverge. The number of events representing the best cost score is not statistically significant (p-value = 0.25).

### 5.1 Panamanian strangler figs and fig wasps have a history of host switching

At broad phylogenetic scales (*e.g.*, sections of figs, genera of fig wasps), there is evidence for significant lineage associations suggesting ecological interactions have manifested themselves in a process of cospeciation (Rønsted et al., 2005; Cruaud et al., 2012). Within the Panamanian community of strangler figs and associated wasps, previous work has identified a breakdown of the one-to-one fig-to-fig wasp association, and has suggested rampant host switching as the process contributing to highly discordant phylogenetic patterns (Molbo et al., 2003; Machado et al., 2005; Jackson et al., 2008). Machado et al. (2005) argue the more appropriate model for the Panamanian community is one in which genetically well-defined fig wasp species coevolve with groups of genetically less well-defined groups of figs. This model argues for a history where pollinator host switching leads to introgression among fig species. Previous studies, however, have drawn inferences within the Panamanian community from limited molecular data and considerable uncertainty in phylogenetic estimates. As limited molecular data can lead to high levels of phylogenetic uncertainty and an inability to recover the true species’ history, our goal was to test these hypotheses harnessing the ability of next-generation sequencing.

Using thousands of RAD loci for the figs and hundreds of UCE loci for the wasps, we provide strong evidence that the history of fig-pollinator association has been punctuated by numerous host switch events in which fig wasps move to and successfully colonize non-natal host lineages through evolutionary time. Host switching is broadly distributed phylogenetically and modeled to have played an important role in shaping the evolutionary association of fig wasps with most of the fig species sampled here (Figure S5). More recent exchanges are also likely to be ongoing and detectable (Cornille et al., 2012). Our results have important implications for evolutionary dynamics within the Panamanian strangler fig community. Host switching provides a mechanism for introgression among fig hosts, which could blur species boundaries between fig species and further lower barriers to host switching in fig pollinators. These effects could span additional trophic levels, reducing host specificity in the diverse community of non-pollinating fig wasp parasites (gallers, kleptoparasites, and parasitoids) and other antagonists associated with each fig species (*e.g.*, Borges, 2015; Marussich and Machado, 2007; Piatscheck et al., 2018; Van Goor et al., 2018; West et al., 1996; Herre, 1993).

Ecological data suggest specificity between fig wasps and their host figs is high, with fig wasps doing a nearly perfect job at locating and pollinating the correct host (Bronstein, 1987, E.A. Herre unpublished). Despite phylogenetic evidence of host switching in the past, our sampling strongly supports current species-specificity, as wasps reared from the same fig species—both from independent fruit on the same tree and from different trees—cluster in phylogenetic space into well-delineated lineages (Figure S3). Our result of species specificity is consistent with previous phylogenetic inference based on mitochondrial DNA in this community (Machado et al., 2005; Molbo et al., 2003). Strong evidence for host-specificity in shallow evolutionary time, combined with host switching in deeper time, focuses attention on the potential ecological and evolutionary processes that could lead to discordant patterns of host association. Floral volatiles released by receptive figs promote species specificity by attracting pollinating wasps carrying conspecific pollen. For example, Proffit et al. (2009) presented the volatile blends of three sympatric fig species to the pollinator of one of those species (*Ficus hispida*). Behavioral assays showed the pollinator is only attracted to its host species, demonstrating host-specificity for this pollinating wasp. Selective pressures on the wasps also play an important role in the maintenance of this interaction. Moe and Weiblen (2012) demonstrated that seed development and growth were comparable for figs pollinated with conspecific and heterospecific pollen in a group of sympatric New Guinea figs, but wasp larval development was reduced for those found in a non-natal host, demonstrating strong selective pressures against wasp development in the incorrect host species. Thus, strong selective pressures on both figs and fig wasps promote species specificity in recent evolutionary time.

Life history traits and geography provide ample opportunity for host shifts to occur and be maintained. Fig wasps have multiple generations per year (8-12) with a large number of pollen-carrying female wasp offspring produced per tree (Weiblen, 2002; Korine et al., 2000). They are capable dispersers, traveling many kilometers in search of a receptive fig (Nason et al., 1996), with some fig wasps having been recorded dispersing over 160 kilometers (Ahmed et al., 2009). Many fig species have large distributions, and in cases where pollinators have been sampled from throughout their range, multiple distinct pollinator species have been found, typically in different geographic areas (*e.g.*, Bain et al., 2016; Chen et al., 2012; Rodriguez et al., 2017). For example, Darwell et al. (2014) explored pollinator species diversity on *Ficus rubiginosa* distributed from northern Queensland to New South Wales in Australia. They discovered the presence of five pollinator species with limited species overlap (although they are not geographically isolated). Style and ovipositor length can overlap across species boundaries (Nefdt and Compton, 1996), allowing for a pollinating wasp to oviposit eggs in a non-associated fig species. Although strong selective pressures drive species specificity, pollinators have been documented in non-natal receptive figs and can produce viable offspring (Ramírez, 1970). In fact, analyses of evolutionary relationships among groups of fig wasp species sharing the same monoecious hosts (copollinators) show that a significant fraction (32%) are not closely related and have thus switched hosts at some point in the past (Yang et al., 2015).

What broader patterns can we infer about figs and fig wasps from results in our local Panama community? In the Neotropics, two sections of figs (strangler figs, *Ficus* subgenus *Urostigma*, section *Americana* and free-standing figs, *Ficus* subgenus *Pharmacosycea,* section *Pharmacosycea*) and their pollinating wasps (*Pegoscapus* and *Tetrapus,* respectively) have independently colonized the region (Cruaud et al., 2012). Strangler figs represent much of the species diversity in the Neotropics, with over 120 described species compared to roughly 20 described species for the free-standing figs. Breakdown of the one-to-one rule and tangled phylogenetic histories of host and symbiont suggests a complex history for strangler figs in the Neotropics. Many of the strangler fig species found in Panama have widespread distributions, and in the broader strangler fig phylogeny, the Panama community is not monophyletic (Machado et al., 2018). It is probable many of these fig species contain multiple pollinator species along their distribution. Within Panama, Molbo et al. (2003) found evidence for four of the eight sampled strangler fig species to contain multiple pollinator species, demonstrating multiple wasp species pollinating the same host in sympatry. Both local sampling through time and geographic sampling of pollinator wasps along the distributions of their host figs will be important for understanding community assembly dynamics of the fig wasps, including where the pollinators came from and how they are related (Molbo et al., 2003; Machado et al., 2005; Jackson et al., 2008; Cornille et al., 2012). We hypothesize that these cases of pollinator sharing and host switching will be mechanistically explainable by similarities of the attractant chemical signals that the receptive host figs produce and the pollinator wasps recognize.

### 5.2 Model-based approaches for addressing cophylogenetic questions

Current cophylogenetic methods measure the degree to which two phylogenies compare (global-fit) or map one tree onto another using *a priori* event costs and the parsimony criterion to identify the optimal reconciliation (event-based). These methods are necessarily limited because they draw inferences based on pattern. As multiple processes can produce similar patterns in the data, results can be challenging to interpret in the absence of an probabilistic approach where the relative contributions of the processes can be statistically evaluated. Thus, it is critical to use a model-based approach to statistically infer the processes important for producing our observed patterns. For example, recent advances in cophylogenetic methods use approximate Bayesian computation (ABC) to estimate the posterior probability of processes important for producing cophylogenetic patterns (Alcala et al., 2017; Baudet et al., 2014). Although ABC is likelihood free and can be sensitive to choice of summary statistics for approximating the posterior probability (Robert et al., 2011), these methods provide an important step forward in evaluating cophylogenetic questions in a process-generating perspective. Our application of the DTL model calculates the likelihood of our data given a model, allowing us to directly measure the evolutionary and ecological processes that have contributed to this obligate mutualism.

Host switching has played a prominent role in shaping the evolution of Panamanian figs and fig wasps. To see how this model-based approach would behave in a system where cospeciation is expected to be the dominant signal, we applied the DTL model to gophers and their chewing lice (see Supporting Information for details; Hafner et al., 1994). Results were drastically different from the figs and fig wasps, and support previous conclusions of cospeciation in this host and parasite system. For gophers and lice, mean maximum likelihood rates were 1.03 (± 0.27) for host switching, 0.72 (± 0.17) for louse extinction, and ~0 for louse speciation (Figure S6). Given these rates and the probabilities of events along the host gopher trees, the reconciled louse tree sets recover a mean average of 14.03 (± 0.93) cospeciations, 6.81 (± 1.18) host switches, 4.53 (± 0.77) louse extinctions, and ~0 louse speciations (Figure S7). Placed in the context of simulations, a dominant signal of cospeciation is strongly supported. Considering host switching and cospeciation, empirical values significantly deviate from their null distributions, with higher cospeciation (p-value=0) and lower host switching (p-value=0) than expected under a model that approximates free host switching (Figure S8). This same pattern is recovered with the maximum likelihood rates, as the empirical estimate for host switching is significantly lower (p-value=0) than the null distribution (Figure S9). As another example of this approach in a coevolutionary system, Groussin et al. (2017) applied the DTL model to mammals and their microbiomes to test whether vertical versus horizontal transmission was the dominant signal in the coevolution of bacterial communities and their hosts. They discovered that 67% of bacterial groups contained a stronger signal of cospeciation than host switching, demonstrating how this model can inform transmission dynamics of microbiome communities. These two examples, in addition to our study on figs and fig wasps, demonstrate the utility of the DTL model for evaluating coevolutionary dynamics using a process-based approach.

Our application of the DTL model provides a promising approach for inferring the contribution of various processes important in the fig and fig wasp mutualism, as well as other systems with strong ecological associations (*e.g.*, hosts and parasites). A logical extension of this approach will be to incorporate absolute divergence time information of the interacting lineages into the model. Currently, the DTL model as applied in ALEml accounts for the relative node positioning within the host phylogeny, but does not account for branch length information of the symbiont trees. While appropriate when addressing gene families within species, as genes evolve within species, this assumption may be violated when dealing with host/symbiont interactions as the timing of association and diversification may not be the same. For example, if diversification times were on different time scales between host and symbiont, some evolutionary processes (*e.g.*, host switching, cospeciation) would effectively not be possible if associated lineages were not present at the same time. Where this would be most problematic is if phylogenetic patterns were similar between interacting lineages, but divergence times at those shared nodes were not overlapping. This pseudocongruence would obscure the true history that a process other than cospeciation necessarily contributed to these patterns (*e.g.*, Donoghue and Moore, 2003). Although there are challenges associated with integrating temporal information in the model-based framework, a more fundamental problem is estimating accurate divergence time information for cophylogenetic analysis. Divergence time estimation is challenging, and timing estimates often contain considerable uncertainty. In addition, without appropriate fossil information, it can be impossible to convert relative branch lengths into absolute time with any sort of certainty. The DTL model provides an approach for measuring the contribution of various processes in the diversification of interacting lineages. It will be important to further adapt the model for cophylogenetic inference, where we can account for additional parameters (*e.g.*, divergence times) important in systems with tight ecological interactions.

## 6 Conclusions

This is the first study to use genome-scale data to address fundamental coevolutionary questions in the fig and fig wasp obligate mutualism. Figs and fig wasps provide a model system for understanding the processes most important for contributing to the maintenance and diversification of an obligate mutualism. Using genomic data coupled with a model-based framework, we demonstrate that host switching has played an important role shaping diversification patterns in the Panamanian strangler fig community. This provides a mechanism for introgression within the figs, a process that has been suggested but not conclusively demonstrated. Adapting a gene-family evolution model for cophylogenetic questions, we provide a framework for evaluating coevolutionary diversification in interacting lineages. Our study highlights the importance of using genome-wide sequence data and a process-based approach for drawing meaningful inferences into the processes structuring and driving the evolution of coevolutionary systems.

## Supporting information

## Acknowledgements

We thank Adalberto Gomez for collecting many of the fig and fig wasp samples. We thank members of the Heath lab, Nason lab, Bastien Boussau, Michael Landis, and April Wright for comments and discussion regarding the manuscript. Funding was provided by the National Science Foundation (DEB-1556853) to JDN, EAH, and TAH.

## References

Ahmed, S., S. G. Compton, R. K. Butlin, and P. M. Gilmartin, 2009. Wind-borne insects mediate directional pollen transfer between desert fig trees 160 kilometers apart. Proceedings of the National Academy of Sciences 106:20342–20347.

Alcala, N., T. Jenkins, P. Christe, and S. Vuilleumier, 2017. Host shift and cospeciation rate estimation from co-phylogenies. Ecology Letters 20:1014–1024.

Bain, A., R. M. Borges, M.-H. Chevallier, H. Vignes, N. Kobmoo, Y. Q. Peng, A. Cruaud, J. Y. Rasplus, F. Kjellberg, and M. Hossaert-Mckey, 2016. Geographic structuring into vicariant species-pairs in a wide-ranging, high-dispersal plant-insect mutualism: the case of Ficus racemosa and its pollinating wasps. Evolutionary Ecology 30:663–684.

Baird, N. A., P. D. Etter, T. S. Atwood, M. C. Currey, A. L. Shiver, Z. A. Lewis, E. U. Selker, W. A. Cresko, and E. A. Johnson, 2008. Rapid SNP discovery and genetic mapping using sequenced RAD markers. PLoS ONE 3:e3376.

Balbuena, J. A., R. Míguez-Lozano, and I. Blasco-Costa, 2013. PACo: a novel procrustes application to cophylogenetic analysis. PLoS ONE 8:e61048.

Baudet, C., B. Donati, B. Sinaimeri, P. Crescenzi, C. Gautier, C. Matias, and M.-F. Sagot, 2014. Cophylogeny reconstruction via an approximate bayesian computation. Systematic Biology 64:416–431.

Bolger, A. M., M. Lohse, and B. Usadel, 2014. Trimmomatic: a flexible trimmer for Illumina sequence data. Bioinformatics 30:2114–2120.

Borges, R. M., 2015. How to be a fig wasp parasite on the fig-fig wasp mutualism. Current Opinion in Insect Science 8:34–40.

Bouckaert, R., J. Heled, D. Kühnert, T. Vaughan, C.-H. Wu, D. Xie, M. A. Suchard, A. Rambaut, and A. J. Drummond, 2014. BEAST 2: a software platform for Bayesian evolutionary analysis. PLoS Computational Biology 10:e1003537.

Branstetter, M. G., B. N. Danforth, J. P. Pitts, B. C. Faircloth, P. S. Ward, M. L. Buffington, M. W. Gates, R. R. Kula, and S. G. Brady, 2017. Phylogenomic insights into the evolution of stinging wasps and the origins of ants and bees. Current Biology 27:1019–1025.

Bronstein, J. L., 1987. Maintenance of species-specificity in a neotropical fig-pollinator wasp mutualism. Oikos 48:39–46.

Bryant, D., R. Bouckaert, J. Felsenstein, N. A. Rosenberg, and A. RoyChoudhury, 2012. Inferring species trees directly from biallelic genetic markers: bypassing gene trees in a full coalescent analysis. Molecular Biology and Evolution 29:1917–1932.

Capella-Gutiérrez, S., J. M. Silla-Martínez, and T. Gabaldón, 2009. trimal: a tool for automated alignment trimming in large-scale phylogenetic analyses. Bioinformatics 25:1972–1973.

Chen, Y., S. G. Compton, M. Liu, and X.-Y. Chen, 2012. Fig trees at the northern limit of their range: the distributions of cryptic pollinators indicate multiple glacial refugia. Molecular Ecology 21:1687–1701.

Chifman, J. and L. Kubatko, 2014. Quartet inference from SNP data under the coalescent model. Bioinformatics 30:3317–3324.

Conow, C., D. Fielder, Y. Ovadia, and R. Libeskind-Hadas, 2010. Jane: a new tool for the cophylogeny reconstruction problem. Algorithms for Molecular Biology 5:16.

Cornille, A., J. Underhill, A. Cruaud, M. Hossaert-McKey, S. Johnson, K. Tolley, F. Kjellberg, S. Van Noort, and M. Proffit, 2012. Floral volatiles, pollinator sharing and diversification in the fig-wasp mutualism: insights from Ficus natalensis, and its two wasp pollinators (South Africa). Proceedings of the Royal Society of London B: Biological Sciences 279:1731–1739.

Crawford, N. G., B. C. Faircloth, J. E. McCormack, R. T. Brumfield, K. Winker, and T. C. Glenn, 2012. More than 1000 ultraconserved elements provide evidence that turtles are the sister group of archosaurs. Biology Letters 8:783–786.

Cruaud, A. and J.-Y. Rasplus, 2016. Testing cospeciation through large-scale cophylogenetic studies. Current Opinion in Insect Science 18:53–59.

Cruaud, A., N. Rønsted, B. Chantarasuwan, L. S. Chou, W. L. Clement, A. Couloux, B. Cousins, G. Genson, R. D. Harrison, P. E. Hanson, et al., 2012. An extreme case of plant-insect codiversification: figs and fig-pollinating wasps. Systematic Biology 61:1029–1047.

Darwell, C. T., S. al Beidh, and J. M. Cook, 2014. Molecular species delimitation of a symbiotic fig-pollinating wasp species complex reveals extreme deviation from reciprocal partner specificity. BMC Evolutionary Biology 14:189.

Donoghue, M. J. and B. R. Moore, 2003. Toward an integrative historical biogeography. Integrative and Comparative Biology 43:261–270.

Doyle, J. and J. Doyle, 1987. Genomic plant DNA preparation from fresh tissue-ctab method. Phytochem Bull 19:11–15.

Eaton, D. A. R., 2014. PyRAD: assembly of de novo RADseq loci for phylogenetic analyses. Bioinformatics 30:1844–1849.

Ehrlich, P. R. and P. H. Raven, 1964. Butterflies and plants: a study in coevolution. Evolution 18:586–608.

Faircloth, B. C., 2015. PHYLUCE is a software package for the analysis of conserved genomic loci. Bioinformatics 32:786–788.

Faircloth, B. C., M. G. Branstetter, N. D. White, and S. G. Brady, 2015. Target enrichment of ultraconserved elements from arthropods provides a genomic perspective on relationships among hymenoptera. Molecular Ecology Resources 15:489–501.

Groussin, M., F. Mazel, J. G. Sanders, C. S. Smillie, S. Lavergne, W. Thuiller, and E. J. Alm, 2017. Unraveling the processes shaping mammalian gut microbiomes over evolutionary time. Nature Communications 8:14319.

Hafner, M. S., P. D. Sudman, F. X. Villablanca, T. A. Spradling, J. W. Demastes, and S. A. Nadler, 1994. Disparate rates of molecular evolution in cospeciating hosts and parasites. Science 265:1087–1090.

Haine, E. R., J. Martin, and J. M. Cook, 2006. Deep mtdna divergences indicate cryptic species in a fig-pollinating wasp. BMC Evolutionary Biology 6:83.

Heled, J. and A. J. Drummond, 2010. Bayesian inference of species trees from multilocus data. Molecular Biology and Evolution 27:570–580.

Herre, E. A., 1993. Population structure and the evolution of virulence in nematode parasites of fig wasps. Science 259:1442–1445.

Herre, E. A., K. C. Jandér, and C. A. Machado, 2008. Evolutionary ecology of figs and their associates: recent progress and outstanding puzzles. Annual Review of Ecology, Evolution, and Systematics 39:439–458.

Hird, S., L. Kubatko, and B. Carstens, 2010. Rapid and accurate species tree estimation for phylogeographic investigations using replicated subsampling. Molecular Phylogenetics and Evolution 57:888–898.

Hutchinson, M. C., E. F. Cagua, J. A. Balbuena, D. B. Stouffer, and T. Poisot, 2017. paco: implementing procrustean approach to cophylogeny in R. Methods in Ecology and Evolution 8:9312–940.

Jackson, A. P., C. A. Machado, N. Robbins, and E. A. Herre, 2008. Multi-locus phylogenetic analysis of neotropical figs does not support co-speciation with the pollinators: the importance of systematic scale in fig/wasp cophylogenetic studies. Symbiosis 45:57–72.

Katoh, K. and D. M. Standley, 2013. MAFFT multiple sequence alignment software version 7: improvements in performance and usability. Molecular Biology and Evolution 30:772–780.

Kissling, W. D., C. Rahbek, and K. Böhning-Gaese, 2007. Food plant diversity as broad-scale determinant of avian frugivore richness. Proceedings of the Royal Society of London B: Biological Sciences 274:799–808.

Korine, C., E. K. Kalko, and E. A. Herre, 2000. Fruit characteristics and factors affecting fruit removal in a panamanian community of strangler figs. Oecologia 123:560–568.

Kubatko, L. S. and J. H. Degnan, 2007. Inconsistency of phylogenetic estimates from concatenated data under coalescence. Systematic Biology 56:17–24.

Lambert, F. R. and A. G. Marshall, 1991. Keystone characteristics of bird-dispersed Ficus in a malaysian lowland rain forest. Journal of Ecology 79:793–809.

Machado, A. F. P., N. Rønsted, S. Bruun-Lund, R. A. S. Pereira, and L. P. de Queiroz, 2018. Atlantic forests to the all Americas: biogeographical history and divergence times of Neotropical Ficus (Moraceae). Molecular Phylogenetics and Evolution 122:46–58.

Machado, C. A., E. Jousselin, F. Kjellberg, S. G. Compton, and E. A. Herre, 2001. Phylogenetic relationships, historical biogeography and character evolution of fig-pollinating wasps. Proceedings of the Royal Society of London B: Biological Sciences 268:685–694.

Machado, C. A., N. Robbins, M. T. P. Gilbert, and E. A. Herre, 2005. Critical review of host specificity and its coevolutionary implications in the fig/fig-wasp mutualism. Proceedings of the National Academy of Sciences 102:6558–6565.

Marussich, W. A. and C. A. Machado, 2007. Host-specificity and coevolution among pollinating and nonpollinating new world fig wasps. Molecular Ecology 16:1925–1946.

McCormack, J. E., B. C. Faircloth, N. G. Crawford, P. A. Gowaty, R. T. Brumfield, and T. C. Glenn, 2012. Ultraconserved elements are novel phylogenomic markers that resolve placental mammal phylogeny when combined with species-tree analysis. Genome Research 22:746–754.

Moe, A. M. and G. D. Weiblen, 2012. Pollinator-mediated reproductive isolation among dioecious fig species (Ficus, Moraceae). Evolution 66:3710–3721.

Molbo, D., C. A. Machado, J. G. Sevenster, L. Keller, and E. A. Herre, 2003. Cryptic species of fig-pollinating wasps: implications for the evolution of the fig-wasp mutualism, sex allocation, and precision of adaptation. Proceedings of the National Academy of Sciences 100:5867–5872.

Nason, J. D., E. A. Herre, and J. L. Hamrick, 1996. Paternity analysis of the breeding structure of strangler fig populations: evidence for substantial long-distance wasp dispersal. Journal of Biogeography 23:501–512.

Nefdt, R. J. C. and S. G. Compton, 1996. Regulation of seed and pollinator production in the fig-fig wasp mutualism. Journal of Animal Ecology 65:170–182.

Piatscheck, F., J. Van Goor, D. D. Houston, and J. D. Nason, 2018. Ecological factors associated with pre-dispersal predation of fig seeds and wasps by fig-specialist lepidopteran larvae. Acta Oecologica 90:151–159.

Proffit, M., C. Chen, C. Soler, J.-M. Bessière, B. Schatz, and M. Hossaert-McKey, 2009. Can chemical signals, responsible for mutualistic partner encounter, promote the specific exploitation of nursery pollination mutualisms?-The case of figs and fig wasps. Entomologia Experimentalis et Applicata 131:46–57.

Rambaut, A., A. J. Drummond, D. Xie, G. Baele, and M. A. Suchard, 2018. Posterior summarization in Bayesian phylogenetics using Tracer 1.7. Systematic Biology 67:901–904.

Ramírez, W., 1970. Host specificity of fig wasps (Agaonidae). Evolution 24:680–691.

Robert, C. P., J.-M. Cornuet, J.-M. Marin, and N. S. Pillai, 2011. Lack of confidence in approximate Bayesian computation model choice. Proceedings of the National Academy of Sciences 108:15112–15117.

Rodriguez, L. J., A. Bain, L.-S. Chou, L. Conchou, A. Cruaud, R. Gonzales, M. Hossaert-McKey, J.-Y. Rasplus, H.-Y. Tzeng, and F. Kjellberg, 2017. Diversification and spatial structuring in the mutualism between Ficus septica and its pollinating wasps in insular South East Asia. BMC Evolutionary Biology 17:207.

Rønsted, N., G. D. Weiblen, J. M. Cook, N. Salamin, C. A. Machado, and V. Savolainen, 2005. 60 million years of co-divergence in the fig-wasp symbiosis. Proceedings of the Royal Society of London B: Biological Sciences 272:2593–2599.

Stadler, T., 2011. Simulating trees with a fixed number of extant species. Systematic Biology 60:676–684.

Starrett, J., S. Derkarabetian, M. Hedin, R. W. Bryson, J. E. McCormack, and B. C. Faircloth, 2017. High phylogenetic utility of an ultraconserved element probe set designed for Arachnida. Molecular Ecology Resources 17:812–823.

Swofford, D. L., 2003. PAUP*: phylogenetic analysis using parsimony (*and other methods). Sinauer Associates, Sunderland, Massachusetts.

Szöllősi, G. J., B. Boussau, S. S. Abby, E. Tannier, and V. Daubin, 2012. Phylogenetic modeling of lateral gene transfer reconstructs the pattern and relative timing of speciations. Proceedings of the National Academy of Sciences 109:17513–17518.

Szöllősi, G. J., W. Rosikiewicz, B. Boussau, E. Tannier, and V. Daubin, 2013a. Efficient exploration of the space of reconciled gene trees. Systematic Biology 62:901–912.

Szöllősi, G. J., E. Tannier, N. Lartillot, and V. Daubin, 2013b. Lateral gene transfer from the dead. Systematic Biology 62:386–397.

Terborgh, J., 1986. Keystone plant resources in the tropical forest. Conservation biology: the source of scarcity and diversity.

Thompson, J. N., 1994. The coevolutionary process. University of Chicago Press.

Van Goor, J., F. Piatscheck, D. D. Houston, and J. D. Nason, 2018. Figs, pollinators, and parasites: A longitudinal study of the effects of nematode infection on fig wasp fitness. Acta Oecologica 90:140–150.

Wang, G., C. H. Cannon, and J. Chen, 2016. Pollinator sharing and gene flow among closely related sympatric dioecious fig taxa. Proceedings of the Royal Society of London B: Biological Sciences 283:20152963.

Weiblen, G. D., 2002. How to be a fig wasp. Annual Review of Entomology 47:299–330.

West, S. A., E. A. Herre, D. M. Windsor, and P. R. Green, 1996. The ecology and evolution of the new world non-pollinating fig wasp communities. Journal of Biogeography 23:447–458.

Wiebes, J. T., 1979. Co-evolution of figs and their insect pollinators. Annual Review of Ecology and Systematics 10:1–12.

Yang, L.-Y., C. A. Machado, X.-D. Dang, Y.-Q. Peng, D.-R. Yang, D.-Y. Zhang, and W.-J. Liao, 2015. The incidence and pattern of copollinator diversification in dioecious and monoecious figs. Evolution 69:294–304.

