## Supplementary material for "Inferring Processes of Coevolutionary Diversification in a Community of Panamanian Strangler Figs and Associated Pollinating Wasps"

### Supporting Information

#### Methods for Gophers and Lice example

We applied the ALEml approach to another empirical system, gophers and their associated lice parasites. This system provides a useful example where the dominant signal is expected to be cospeciation, due to the evolutionary tracking of parasites with their gopher hosts. We downloaded the COI mitochondrial DNA sequence data sets from Hafner et al. (1994), composed of 15 taxa for the gophers and 17 taxa for the lice, and used these data as outlined in the framework above. Briefly, sequences were aligned with muscle v3.8.31 (Edgar, 2004), and gene tree distributions estimated in BEAST v2.4.7. Both data sets used a birth-death model as a tree prior, a TrN + I + $\Gamma$ substitution model, and a strict clock model. Analyses were run for 100 million generations, sampling every 10,000 generations, which yielded 10,000 samples from the target distribution; convergence was assessed with Tracer. We used the ALE pipeline as outlined above, using the posterior distribution of gene trees for the lice as the "gene trees" and the posterior distribution of gene trees for the gophers as the "species trees", integrating over 1,000 trees from the gopher tree distribution to account for phylogenetic uncertainty. Simulations were conducted to construct null distributions for the model parameters, simulating 1,000 trees to match the louse sampling (as described above for the fig and fig wasp system). For ALE simulations, we used the gopher MCC BEAST tree while analyzing 1,000 simulated trees. Parameter estimates were summarized as null distributions with which to compare empirical estimates.

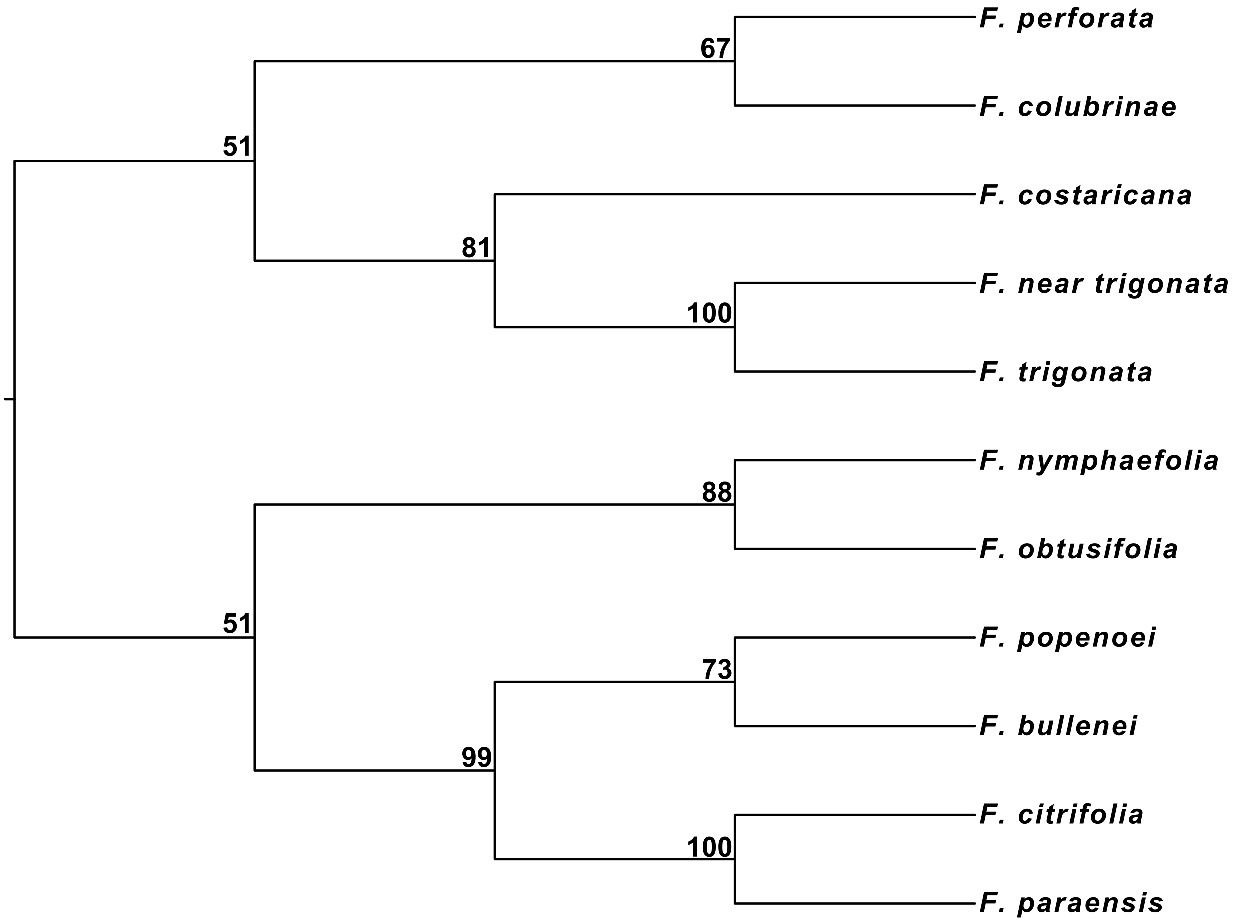


**Figure S1: SVDQ$_{ST}$ of the figs.** Rooting was informed by results from the SNAPP analysis.

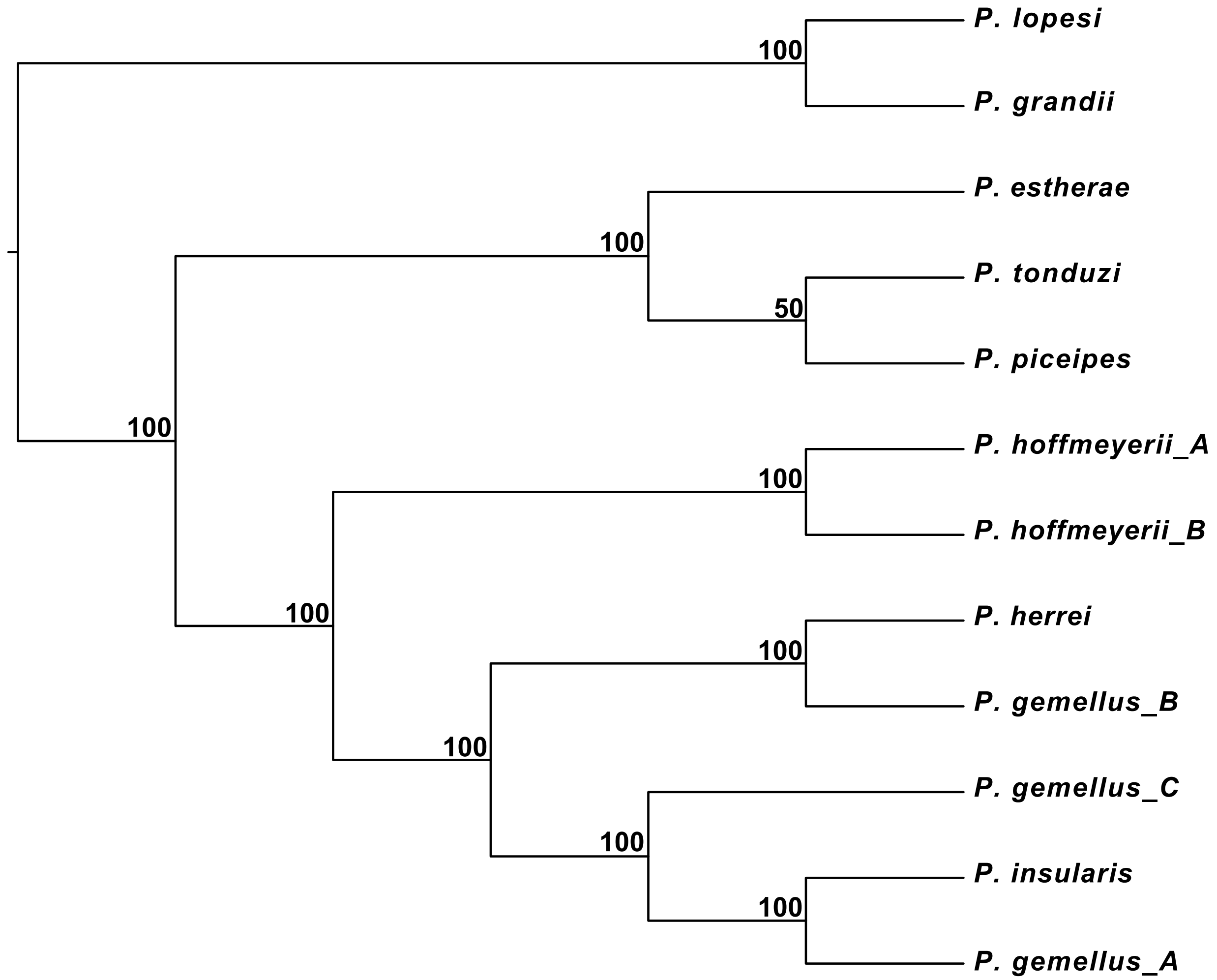


**Figure S2: SVDQ$_{ST}$ of the fig wasps.** Rooting was informed by results from the StarBEAST2 analysis.

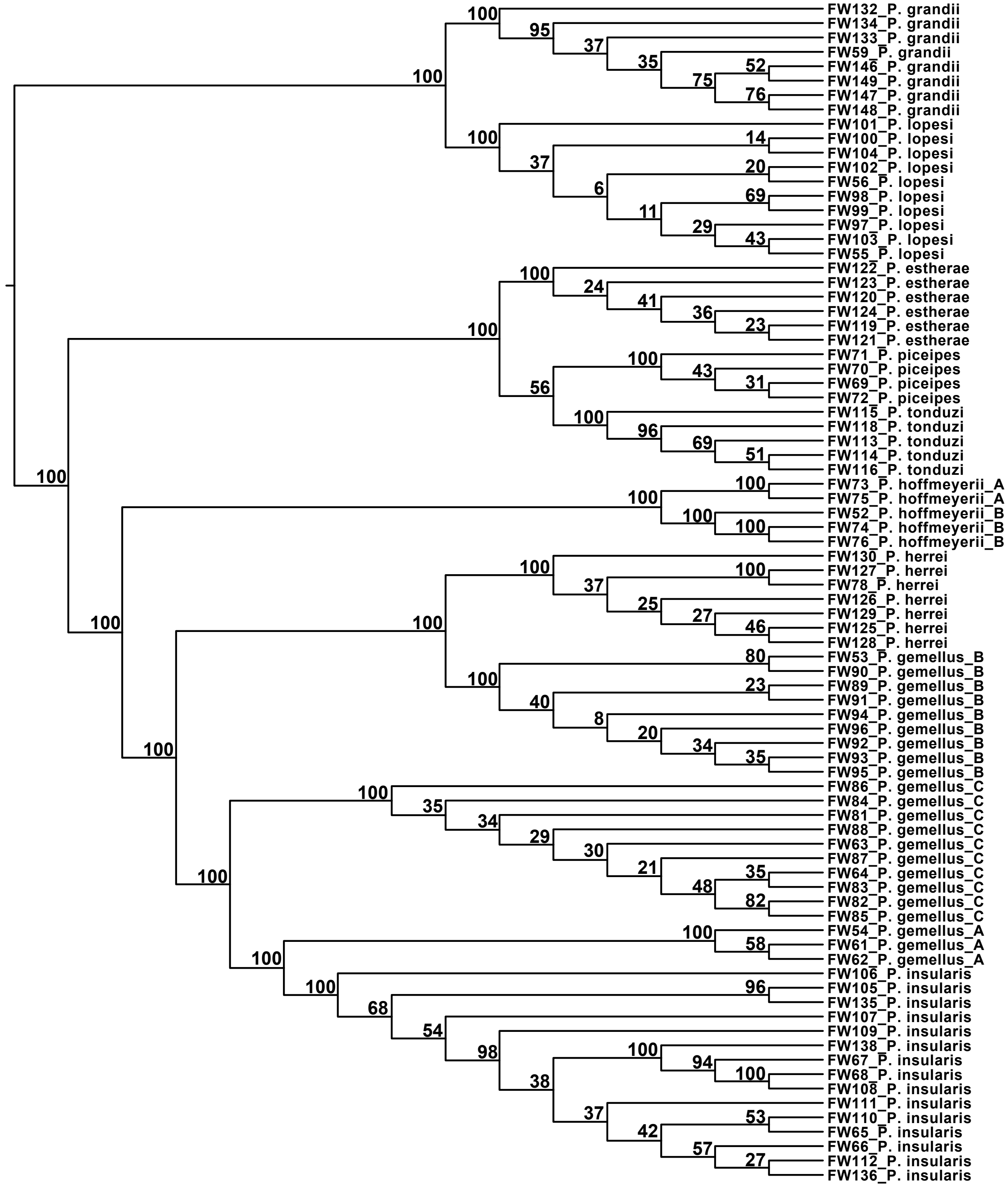


**Figure S3: SVDQ$_{LT}$ of the fig wasps.** Rooting was informed by results from the StarBEAST2 analysis.

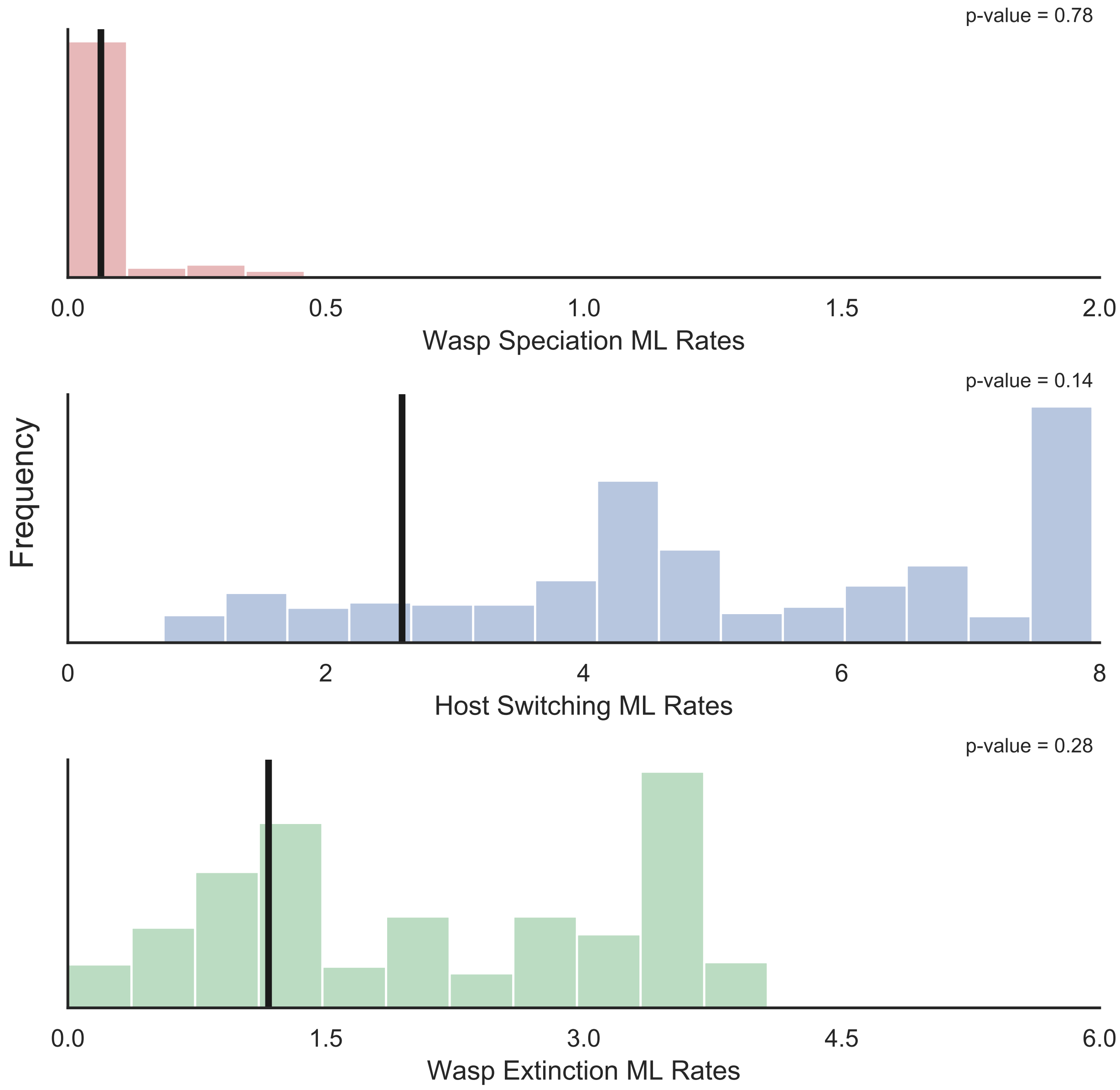


**Figure S4: Maximum likelihood rates when using the fig MCC species tree and a simulated fig wasp species tree.** Null distributions are based on 953 simulated fig wasp species trees (some ALEml analyses returned an error). The vertical line represents the mean estimate from the empirical data. For wasp speciation, host switching, and wasp extinction, p-values represent the proportion of simulations that produced an estimate equal to or below the empirical value.

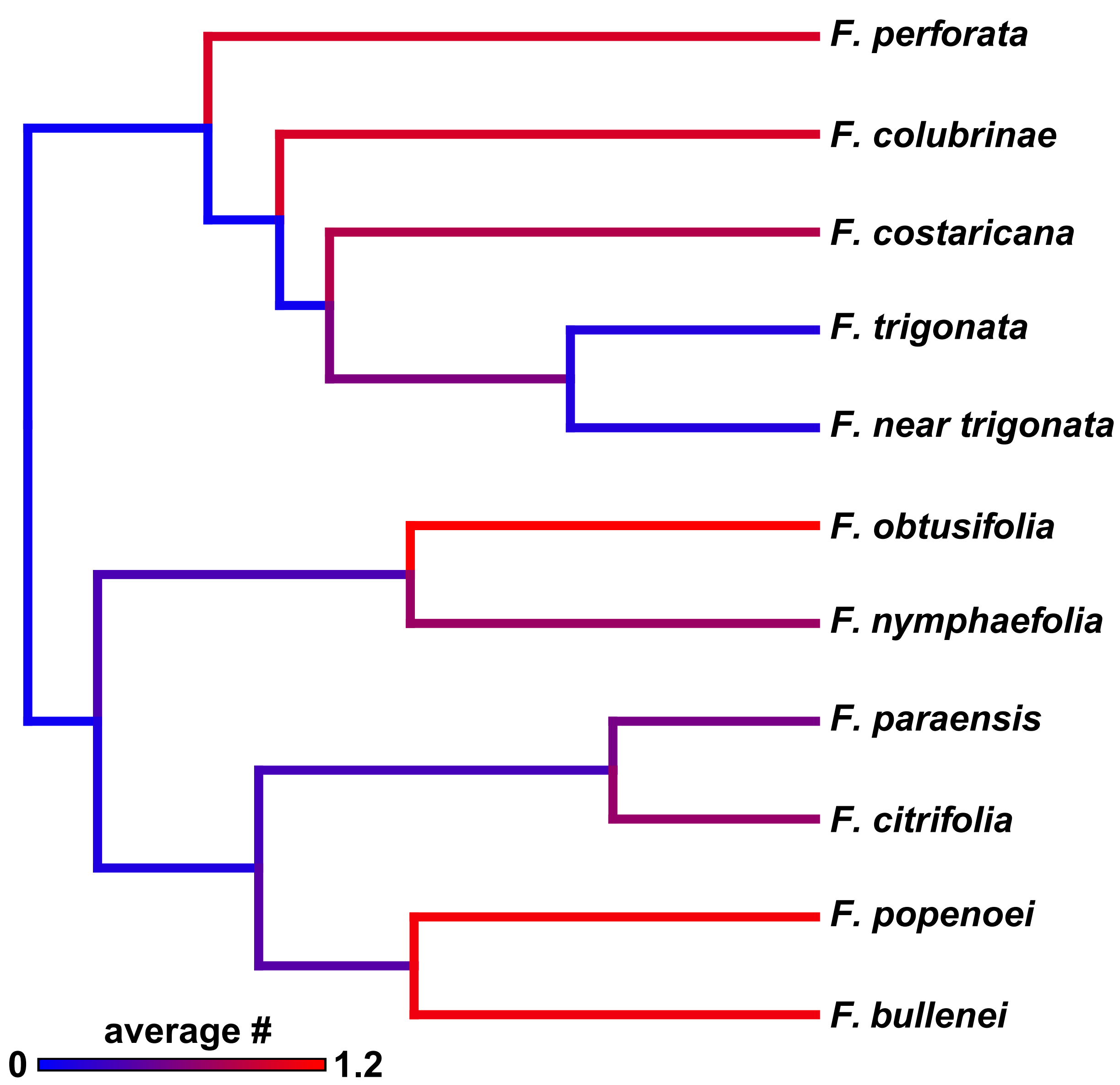


**Figure S5: Average number of host switching events for the figs and fig wasps.** The MCC species tree representing the figs (SNAPP) was used as the host tree in the ALEml analysis. Host switching events are represented among the branches of the tree.

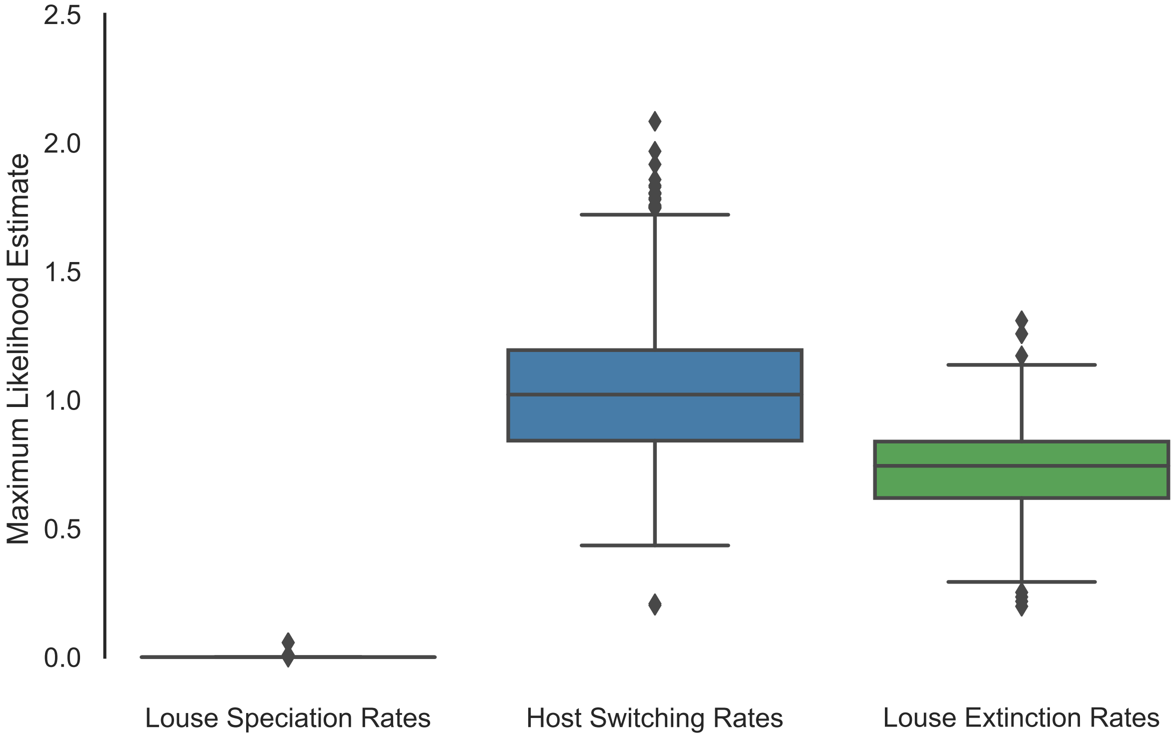


**Figure S6: Maximum likelihood estimates for the rates of the three processes estimated in ALEml for the analysis of the gopher/louse host-parasite system.** Results are averaged across 1,000 trees drawn from the posterior distribution of species trees representing the gophers.

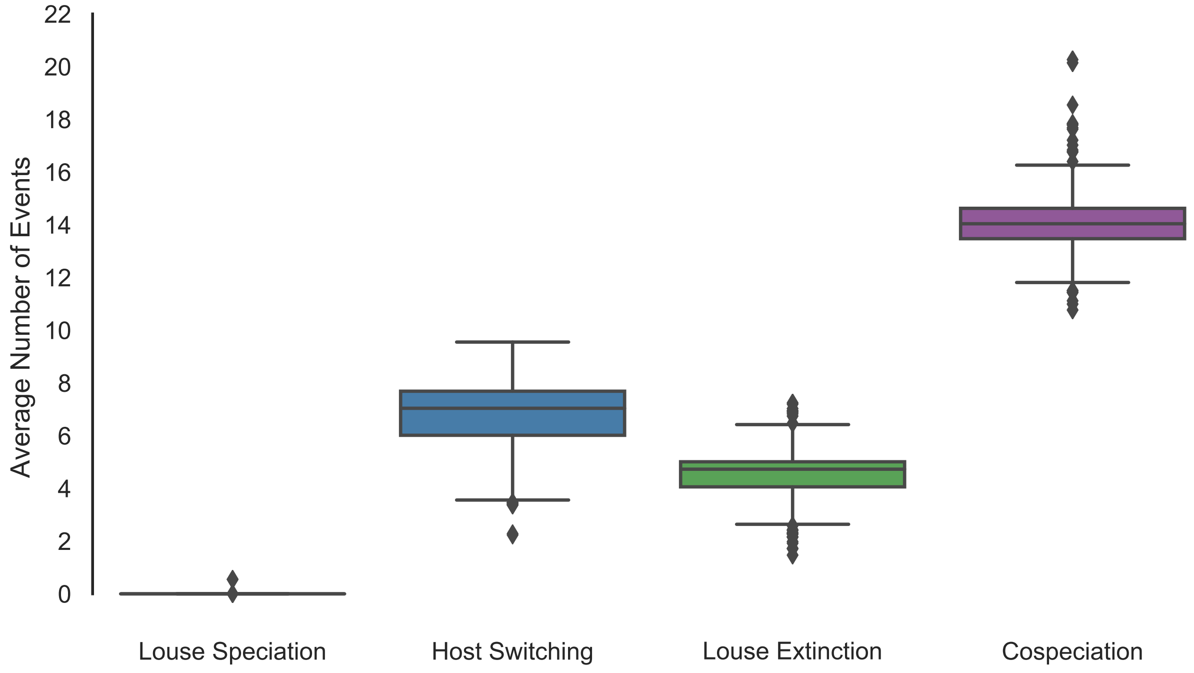


**Figure S7: Average number of events estimated from ALEml.** Results are averaged across 1,000 trees drawn from the posterior distribution of species trees representing the gophers.

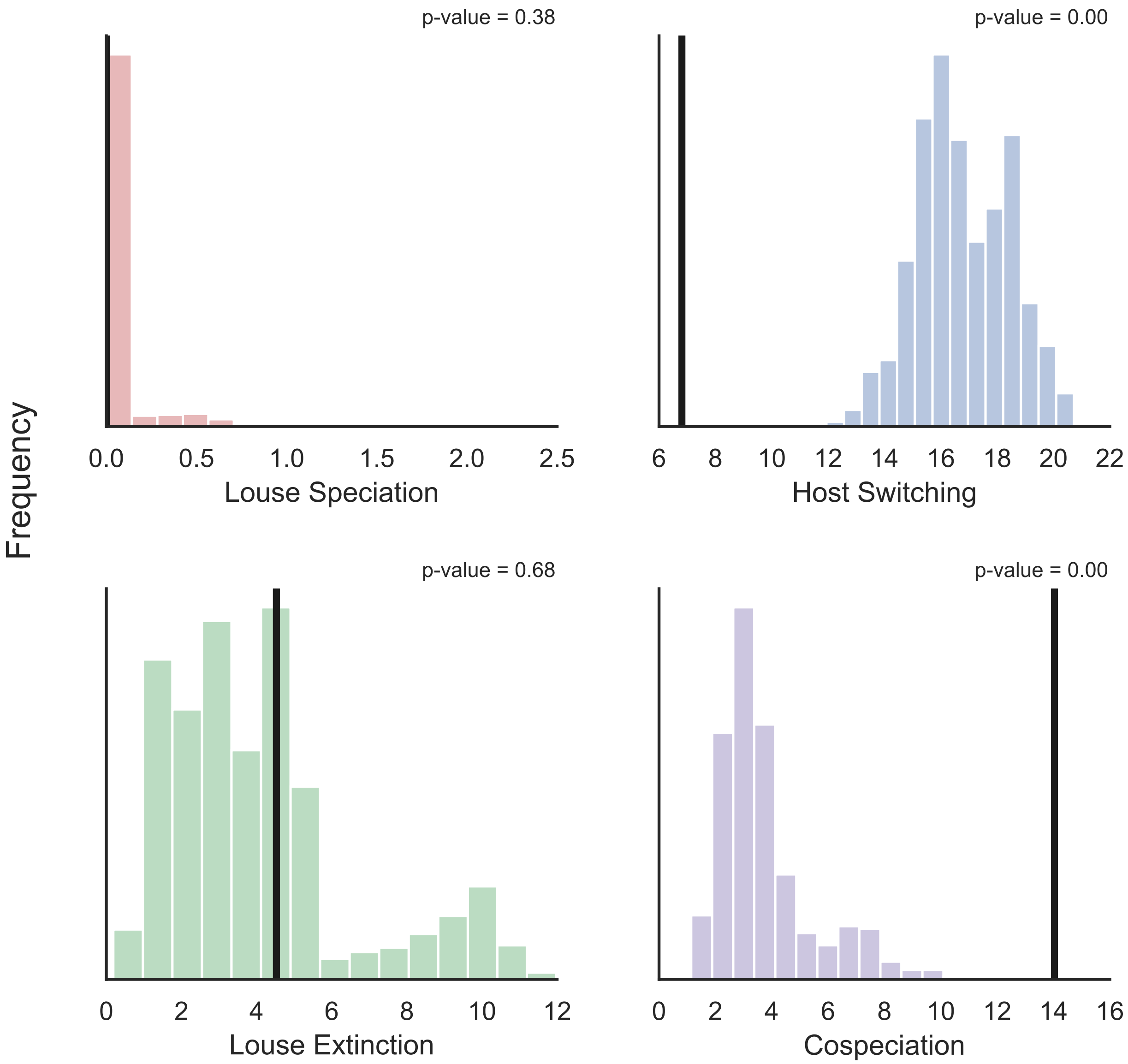


**Figure S8: Number of events when using the gopher MCC species tree and a simulated louse species tree.** Null distributions are based on 932 simulated louse species trees (some ALEml analyses returned an error and were subsequently skipped). The vertical line represents the estimate from the empirical data. For louse speciation, louse extinction, and host switching, p-values represent the proportion of simulations that produced an estimate equal to or below the empirical value. For cospeciation, the p-value (0.00) represents the proportion of simulations that produced an estimate equal to or above the empirical value.

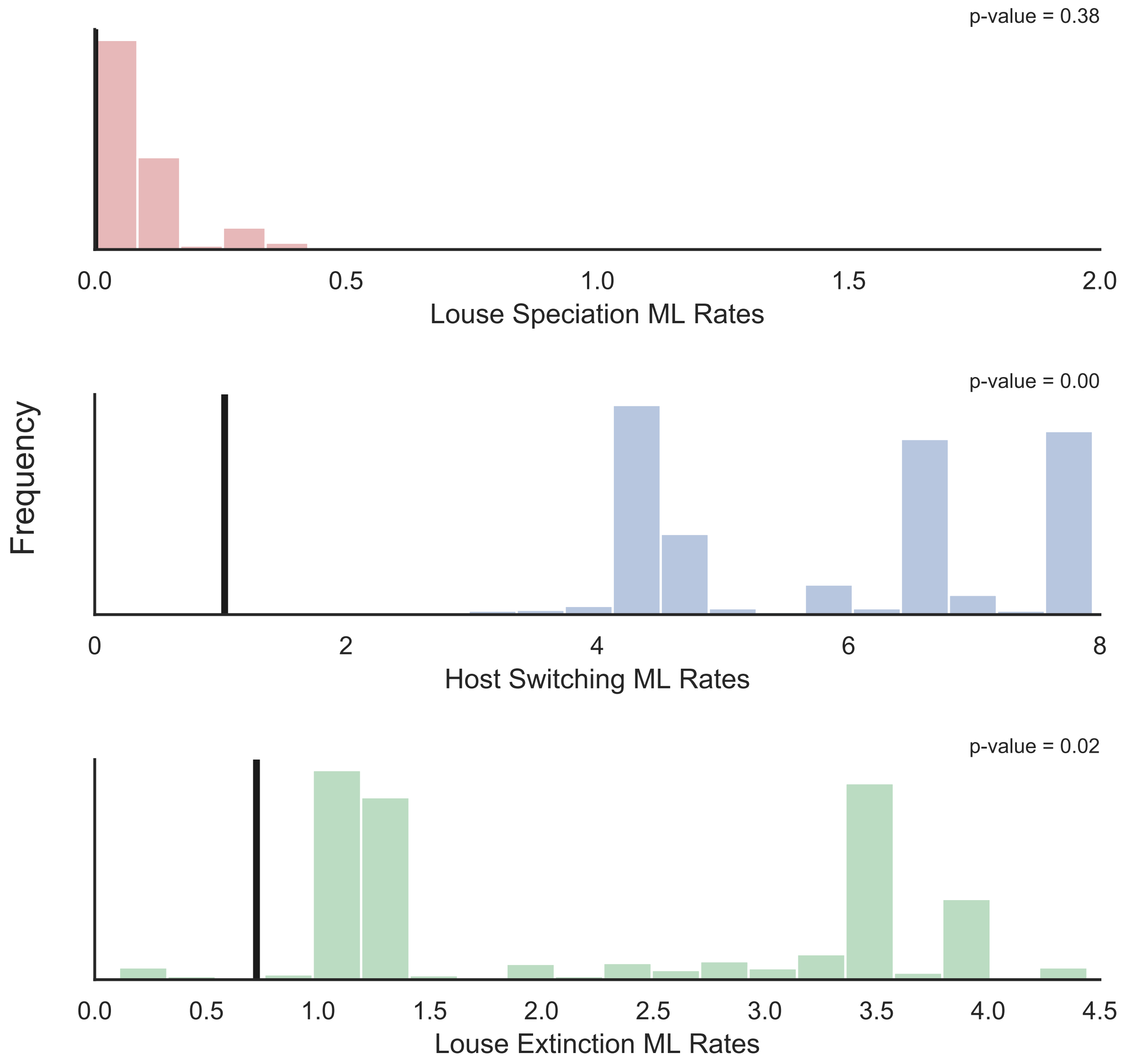


**Figure S9: Maximum Likelihood rates when using the gopher MCC species tree and a simulated louse species tree.** Null distributions are based on 932 simulated louse species trees (some ALEml analyses returned an error and were subsequently skipped). The vertical line represents the mean estimate from the empirical data. For louse speciation, host switching, and louse extinction, p-values represent the proportion of simulations that produced an estimate equal to or below the empirical value.

### References

Edgar, R. C., 2004. Muscle: multiple sequence alignment with high accuracy and high throughput. Nucleic Acids Research 32:1792–1797.

Hafner, M. S., P. D. Sudman, F. X. Villablanca, T. A. Spradling, J. W. Demastes, and S. A. Nadler, 1994. Disparate rates of molecular evolution in cospeciating hosts and parasites. Science 265:1087–1090.
